# Synaptic Proteome Alterations in the Primary Auditory Cortex of Schizophrenia

**DOI:** 10.1101/639914

**Authors:** Matthew L. MacDonald, Megan Garver, Jason Newman, Zhe Sun, Joseph Kannarkat, Ryan Salisbury, Jill Glausier, Ying Ding, David A. Lewis, Nathan Yates, Robert A. Sweet

**Author notes:** Corresponding Author: Matthew L MacDonald, PhD, Assistant Professor of Psychiatry, Translational Neuroscience Program, BioMedical Mass Spectrometry Center, Thomas E. Starzl Biomedical Science Tower, Rm W1655, 203 Lothrop St, Pittsburgh, PA 15213.

## Abstract

**Importance:** Findings from unbiased genetic studies have consistently implicated synaptic protein networks in Schizophrenia (**Sz**), but the molecular pathology at these networks and their potential contribution to the synaptic and circuit deficits thought to underlie disease symptoms remain unknown.

**Objective:** To determine if protein levels are altered within synapses from primary auditory cortex (**A1**) of subjects with Sz; and if so, are these differences restricted to the synapse or present throughout the grey matter?

**Design:** A paired case-control design was utilized for this study. Biochemical fractional – targeted Mass Spectrometry (**MS**) was used to measure the levels of >350 proteins in A1 grey matter homogenate and synaptosome preparations, respectively. All experimenters were blinded to diagnosis at every stage of sample preparation, MS analysis, and raw data processing. The effects of postmortem interval (**PMI**) and antipsychotic drug treatment on protein levels were assessed in mouse and monkey models, respectively.

**Setting:** All cases were recruited from a single site, The Allegheny County Office of the Medical Examiner, and all tissues were processed at the University of Pittsburgh.

**Participants:** Brain specimens from all subjects were obtained during autopsies conducted at the Allegheny County Office of the Medical Examiner after receiving consent from the next-of-kin. An independent panel of experienced clinicians made consensus Diagnostic and Statistical Manual of Mental Disorders Fourth Edition diagnoses. Unaffected comparison subjects underwent identical assessments and were determined to be free of lifetime psychiatric illness. Each Sz subject was matched by sex, and as closely as possible for age and PMI, with one unaffected comparison subject.

**Main Outcomes and Measures:** Primary measures were homogenate and synaptosome protein levels and their co-regulation network features. Prior to data collection we hypothesized: **1.** That levels of canonical postsynaptic proteins in A1 synaptosome preparations would differ between Sz and control subjects; and **2.** That these differences would not be explained by changes in total A1 homogenate protein levels.

**Results:** Mean subject age was 48 years for both groups with a range of 17-83; each group included 35 males and 13 females; mean PMI was 17.7 hours in controls and 17.9 in Sz. We observed robust alterations (q < 0.05) in synaptosome levels of canonical mitochondrial and postsynaptic proteins that were highly co-regulated and not readily explained by postmortem interval, antipsychotic drug treatment, synaptosome yield, or underlying alterations in homogenate protein levels.

**Conclusions and Relevance:** Our findings indicate a robust and highly coordinated rearrangement of the synaptic proteome likely driven by aberrant synaptic, not cell-wide, proteostasis. In line with unbiased genetic findings, our results identified alterations in synaptic levels of postsynaptic proteins, providing a road map to identify the specific cells and circuits that are impaired in Sz A1.

## Introduction

Subjects with schizophrenia (**Sz**) display impairments in the processing of auditory sensory information (1-4) including auditory event-related potentials (e.g. mismatch negativity)(1, 2, 5, 6) which are generated in layer 3 of primary auditory cortex (**A1**) (7-9). Many individuals with Sz also have deficits in auditory learning (10), limiting their functional recovery during targeted sensory and cognitive training(11). In A1, as in other cortical regions, learning requires the formation and stabilization of new dendritic spines (12-14) and decreased density of layer 3 dendritic spines has been reproducibly observed in multiple brain regions in Sz, including A1 (15-20).

Unbiased genetic studies of Sz have implicated synaptic protein networks in disease etiology (21). This convergence of genetic risk factors is believed to underlie the dendritic spine deficits and information processing alterations observed in the A1 and other cortical regions. Numerous studies have documented the essential role of local synaptic protein trafficking, translation, and degradation in spine formation and plasticity (22). This local protein homeostasis (proteostasis) is believed to be required for the rapid synaptogenesis and plasticity essential to cortical information processing and new learning(23, 24). Indeed, blocking synaptic proteostatic processes impairs new synapse formation and learning(25, 26). Thus, we hypothesized that protein levels at the synapse are altered in the A1 of Sz and that these synaptic alterations are distinct from protein alterations in total grey matter.

To investigate protein levels at the synapse, we utilized a high precision targeted mass spectrometry (**MS**) approach with a [^13^C_6_]lysine-labeled brain proteome internal standard ([^**13**^**C**_**6**_**]brain ISTD**)(27) to quantify >350 selected proteins in A1 synaptosome enrichments (to measure protein levels within synapses) from 48 pairs of Sz and unaffected comparison subjects, matched for age, sex, and PMI. While targeted-MS is limited in the number of proteins assayable, it has exceptional precision, accuracy, and throughput. To determine if the predicted alterations are synaptosome-specific, identical studies were conducted in total A1 homogenate preparations to measure tissue-wide protein levels. Technical controls were utilized to monitor reproducibility of synaptosome enrichment, sample preparation, and instrument performance. The effects of antipsychotic drug (**APD**) treatment and postmortem interval (**PMI**) were assessed in monkey and mouse models, respectively.

We observed robust and highly co-regulated differences in the synaptic levels of canonical mitochondrial and postsynaptic proteins between subject groups, including both glutamate and GABA receptors. Tissue-wide levels of a majority of these proteins did not differ significantly in Sz. These findings indicate a broad and highly coordinated rearrangement of the synaptic proteome likely contributing to synapse and circuit pathology. That these synaptic proteome alterations were largely unexplained by tissue-wide protein level differences suggests they are driven by impairments in synaptic, not cell-wide, proteostasis.

## Materials & Methods

**Figure 1** provides an overview the experimental design.

**Figure 1.**
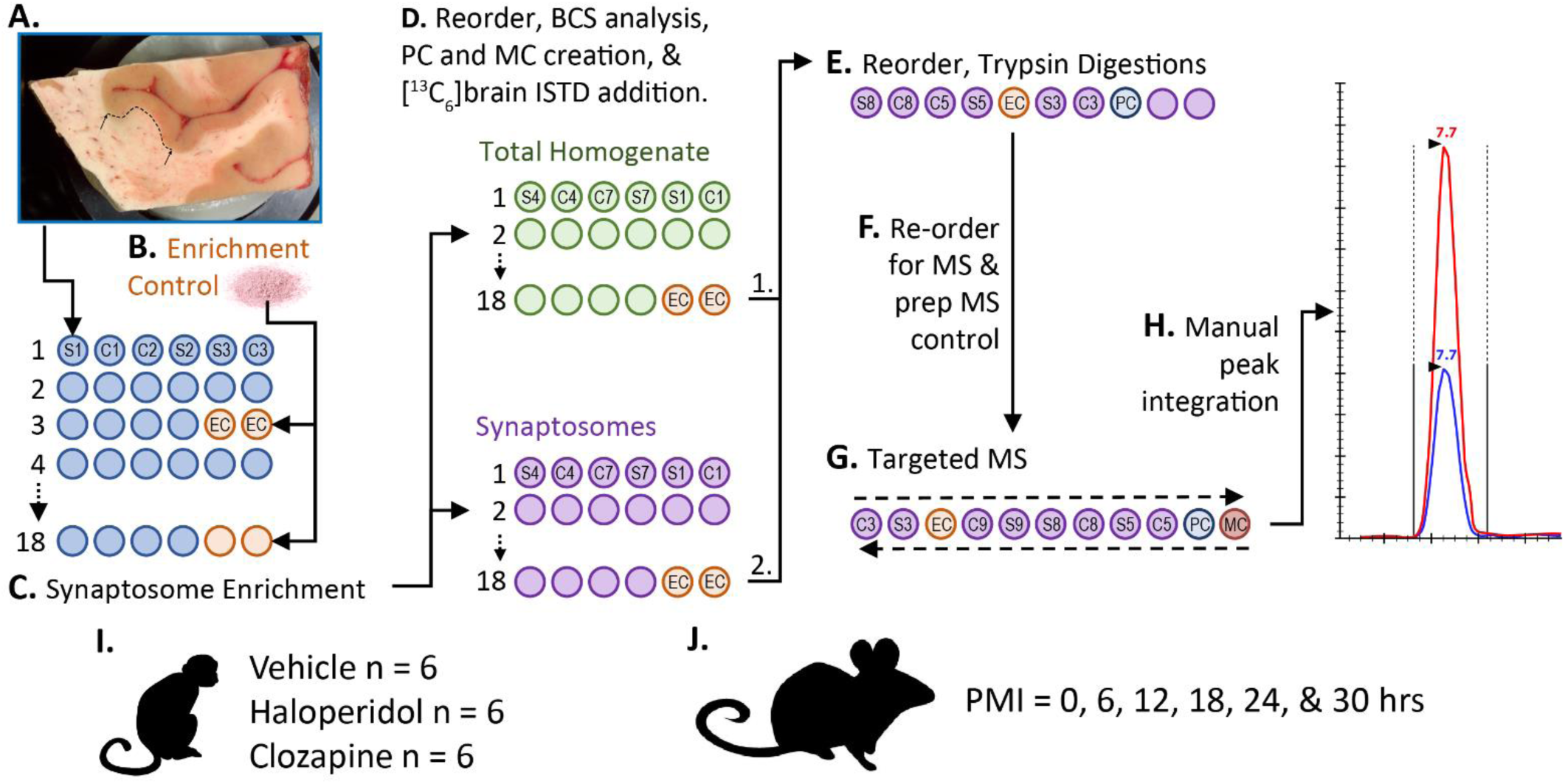
Study Overview. Each Sz subject was paired with a control (matched for sex and as closely as possible for age and PMI, **Table S1**). Paired subject samples were processed together throughout all experiment steps (e.g. S1 & C1, S2 & C2). A randomized block design was utilized with different sample orders for tissue collection, synaptosome enrichment, standard addition, trypsin digestion, and targeted MS. Methodological controls were included to monitor variably in synaptosome enrichment; standard addition and trypsin digestion; and targeted MS. All experimenters were blinded during sample preparation, MS analysis, and peak integration. **A.** Grey matter was collected from Heschl’s Gyrus in 40 μm sections and frozen at −80°C. **B.** The ultra-centrifuge rotor used for synaptosome enrichment has limited capacity at 6 tubes. To monitor reproducibility across synaptosome enrichment 1g of superior temporal gyrus grey matter from a low PMI control was pulverized in liquid nitrogen and divided into 50 mg aliquots. **C**. Two aliquots of these Enrichment Controls (**EC**) were included in every third rotor run. **D.** The resulting homogenate and synaptosome fractions were then reordered into separate blocks and processed independently for the remainder of the protocol. Micro-BCA was used to determine total protein amounts. A pooled control (**PC**) was then created from all samples within each fraction (homogenate or synaptosome) and divided into 8 aliquots to monitor variability in addition of [^13^C_6_]brain ISTD and trypsin digestion. **E.** The samples were again reordered and digested with trypsin. **F.** The samples were reordered a final time, and a single mass spectrometry control (**MC**) to monitor instrument stability was created by mixing a homogenate PC and a synaptosome PC. **G.** All samples were analyzed in duplicate, with the sample order run first forwards and then backwards. ECs were distributed randomly throughout the run order, while PCs and the MC were analyzed every 10-12 injections. **H.** Integration of all light/heavy peak pairs were manually checked, and re-integrated if needed, by trained personnel blinded to sample. **I.** Superior temporal gyrus grey matter from male monkeys treated chronically with haloperidol, clozapine, or vehicle (n = 6/group) was used to monitor antipsychotic drug effects. **J.** Cortical hemispheres from mice with modeled PMIs from 0 - 30 hours were used to monitor the effects of PMI on homogenate and synaptosome protein levels. Monkey grey matter and mouse cortices were processed and analyzed contemporaneously and using an identical protocol and [^13^C_6_]brain ISTD preparation as described above, with the omission of the ECs.

### Human Subjects and Tissue Collection

Brain specimens were obtained from the Allegheny County Office of the Medical Examiner (**Table 1**) and grey matter was harvested from the auditory cortex as previously described(28, 29). See *Supplement* and **Figure 1** for details.

**Table 1.** Summary of Subject Characteristics. There were no diagnostic group differences in age, sex, postmortem interval, storage time, or in the distribution of handedness between the diagnostic groups. There was a small, but significant, decrease in pH in Sz (p = 0.012). *A, ambidextrous; F, female; L, left-handed; M, male; PMI, postmortem interval; R, right-handed; U, unknown.*

|  | Control | Schizophrenia |
| --- | --- | --- |
| N | 48 | 48 |
| Mean Age (SD) | 48.6 (13.5) | 48.1 (13.6) |
| Range | 17-82 | 17-83 |
| Sex (F/M) | 13/35 | 13/35 |
| Handedness (R/L/A/U) | 30/8/1/9 | 42/4/0/2 |
| PMI (SD) | 17.7 (5.7) | 17.9 (7.1) |
| pH (SD) | 6.70 (0.27) | 6.55 (0.31) |
| Storage Time Months (SD) | 155.4 (50) | 148.7(54) |
| Suicide, N (%) | 0 (0%) | 12 (25%) |
| Schizoaffective, N (%) | NA | 14 (29%) |
| Alcohol/Substance abuse ATOD, N (%) | 18 (37.5%) | 29 (60%) |
| Antipsychotic ATOD, N (%) | 0 (0%) | 42 (86.5%) |

### Antipsychotic Drug Treated Monkey Tissue

The tissue utilized here was obtained from previously completed experiments that have been extensively described elsewhere(30). See *Supplement* for details.

### Mouse PMI Brain Tissue

C57BL/6 Adult Mice, n= 2 per timepoint (0, 6, 12, 18, 24, and 30 hrs), were sacrificed by CO_2_ asphyxiation followed by cervical dislocation. The carcasses were incubated at room temperature for 2/3 the PMI duration and then placed at 4°C for the final 1/3. The brain was then removed from the skull, the cerebellum discarded, and the hemispheres separated (giving n = 4 hemispheres/time point), flash frozen in isopentane on dry ice, and stored at −80°C.

### Sample Preparation

Total grey matter homogenate and synaptosome preparations were obtained using a variation on our sucrose density gradient centrifugation method, validated for use in human postmortem brain tissue (27, 31). The homogenate preparation is composed of all material present in the A1 grey matter, an aliquot of which is set aside prior to synaptosome enrichment. See *Supplement* for detailed protocol. To monitor variability in synaptosome preparation in the human cohorts, two 50 mg aliquots of pulverized superior temporal gyrus grey matter from a low PMI human control were included in every third run and assayed alongside subject samples (**Figure 1**). To monitory variability in sample preparation after synaptosome enrichment, a pooled control was prepared and assayed alongside subject samples (**Figure 1**).

### Sample Preparation of Targeted-MS

10 μg homogenate, synaptosome, or pooled controls were mixed with 10 μg of [^13^C_6_]brain ISTD (27) prepared from stable isotope labeling in mammals (SILAM) mouse cortex tissue (Cambridge Isotopes). The human - [^13^C_6_]brain ISTD mixtures were subject to trypsin digestion by FASP(32).

### Development and Validation of Peptide Selected Reaction Monitoring (SRM) Method

Peptide SRMs were generated using data from discovery MS analyses of human grey matter homogenates as well as synaptosome and postsynaptic density enrichments and validated as previously described (27).

### Mass Spectrometry

MS analyses were conducted on a TSQ Quantiva triple stage quadrupole mass spectrometer (ThermoFisher Scientific, Location) with an Ultimate 3000 HPLC (Dionex). 2 µl (∼1 μg protein) was loaded on to a 3 µm 120A; 105mm REPROSIL-Pur C18 Picochip (New Objective) at 1 μl/min for 12 min and eluted at 400nl/min over a 25 min gradient from 3-35% mobile phase B (Acetonitrile, 0.1% formic acid). Selected reaction monitoring transitions were scheduled with 45 second windows. Transitions were monitored, allowing for a cycle time of 1.1 sec, resulting in a dynamic dwell time never falling below 10 msec. The MS instrument parameters were as follows: capillary temperature 275°C, spray voltage 1350 V, and a collision gas of 1.4 mTorr (argon). The resolving power of the instrument was set to 0.7 Da (Full Width Half Maximum) for the first and third quadrupole. Data were acquired using a Chrom Filter peak width of 4.0 sec.

### Data processing

Peak areas and area ratios were calculated within Skyline (33). All individual selected reaction monitoring transitions and integration areas were manually inspected and re-integrated as needed by trained experimenters.

### Statistics

*Reproducibility of Peptide Quantification and Calculation of Protein Level Measures:* See *Supplement*.

#### Proteins enriched in synaptosomes

Of the 348 proteins quantified in the synaptosome fractions, 159 had synaptosome levels that were significantly enriched over their homogenate levels as defined by a Bonferroni corrected p < 0.05 and a synaptosome/homogenate fold change > 1.25 (**Table S4**). Statistical analysis of Sz-control synaptosome differences were limited to these proteins.

#### Effects of PMI

See *Supplement*

#### Differential Expression Analysis

Limma-Voom(34) was performed to detect the difference between Sz and control samples in homogenate and synaptosome proteins, respectively. Both unadjusted and adjusted p-values were obtained (**Tables S2 and S3**). For additional detail see *Supplement*.

#### Network Analysis

Weighted Gene Co-expression Network Analysis (WGCNA)(35) was used to investigate the pattern of co-regulated protein expression. See *Supplement* for details.

#### Comparison to PsychENCODE transcriptome findings

See *Supplement*.

Tables **S2** and **S3** report the module memberships for homogenate and synaptosome proteins, respectively. Tables **S9** and **S10** report the Eigenprotein values for each subject from the homogenate and synaptosome modules. Tables **S11** and **S12** report the full list of enriched terms for the homogenate and synaptosome modules, respectively.

## Results

### Synaptosome protein yield is lower in Sz A1

Total synaptosome yield (ug protein / mg grey matter) was 33% lower (paired t-test, −0.21, 95% CI: −0.31 to −0.1, p = 4E^-4^) in Sz subjects (**Figure 2A**). Haloperidol-treated monkeys had ∼30% greater synaptosome yield relative to vehicle (unpaired t-test, 0.25±0.8, 95% CI: 0.047 to 0.44, p = 0.02, **Figure 2B**). PMI was associated with decreased synaptosome yield in the mouse PMI model (r = −0.84, CI = −0.98 to −0.1, p = 0.035, **Figure 2C**), but not in the human cohort (**Figure S1A**). pH correlated with synaptosome yield in the human cohort (r = 0.37, CI = 0.1746 to 0.5418, p = 4E^-4^, **Figure S1B**); however, yield was still significantly decreased in Sz (linear regression, p = 0.011) while controlling for pH. To control for the lower synaptosome yield in Sz, the same amount of total synaptosome protein from all subjects and animals was used for MS analyses.

**Figure 2.**
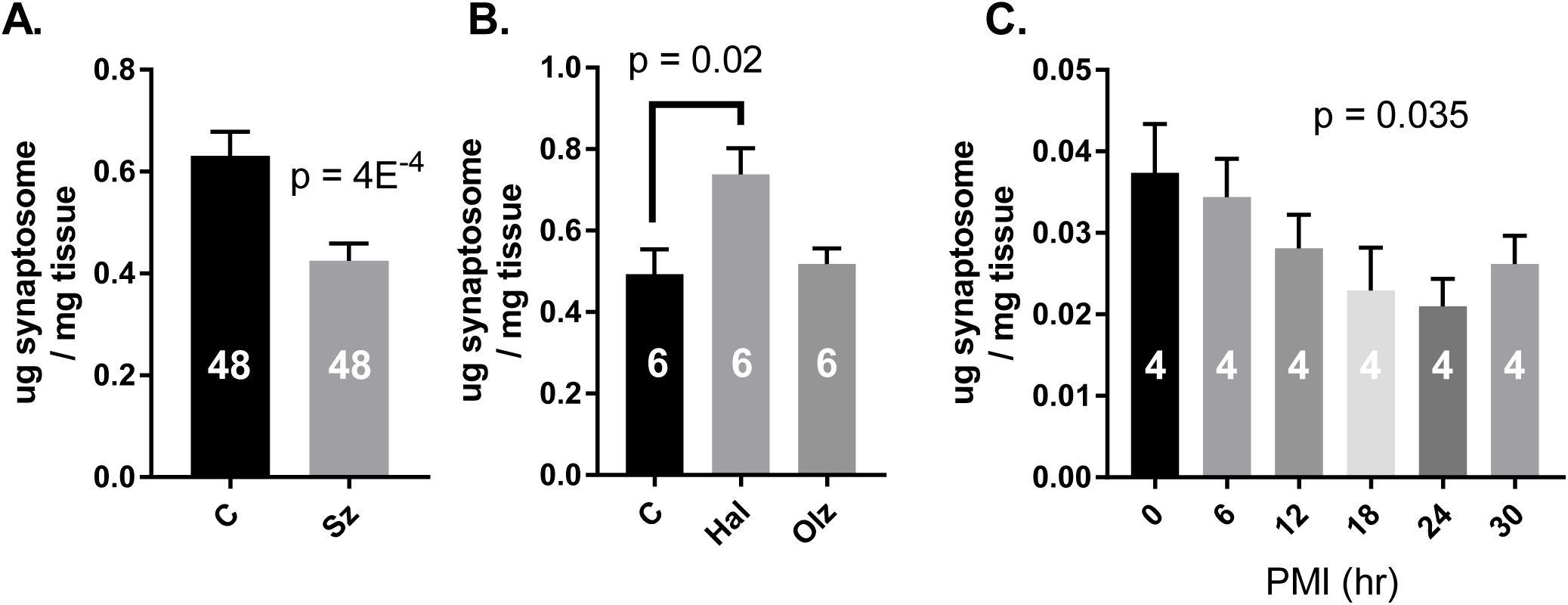
Synaptosome Yield. Total µg of protein in each synaptosome preparation was measured by micro-BCA. This value was then normalized to wet tissue weight to calculate synaptosome yield (µg total synaptosome protein / mg tissue). Synaptosome yield was decreased in Sz (**A.**) and increased in superior temporal gyrus tissue from monkeys treated with haloperidol, but not olanzapine (**B.**). Synaptosome yield was decreased in a linear fashion with PMI in the mouse cortical tissue (**C.**). Group n is provided in each bar.

### Homogenate and synaptosome proteome alterations in Sz

55/402 (14%) proteins in homogenates and 64/155 (41%) in synaptosomes differed significantly between Sz and control subjects (q < 0.05, **Figure 3A & B**). Of the 64 proteins altered in synaptosomes, only 26 could be explained by concurrent changes in homogenate levels of these proteins (**Figure 3C & D**).

**Figure 3.**
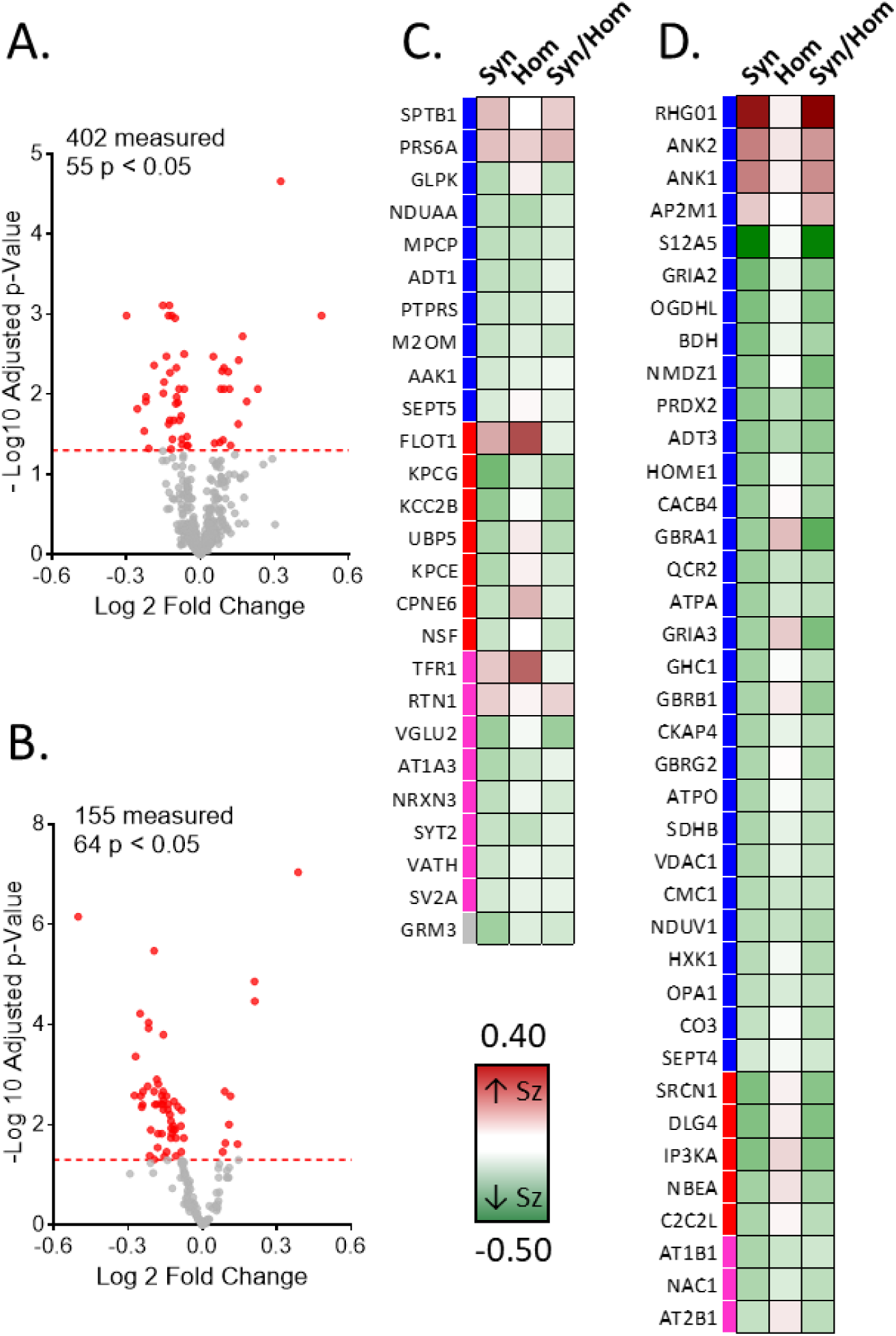
Volcano Plots and Heat Maps of Sz-control Differences. A & B. These volcano plots chart the - log 10 p-value and the log 2-fold change (Sz vs control) for homogenate (**A.**) and synaptosome protein (**B.**) levels. The red dashed lines are set at an adjusted Limma p = 0.05. **C & D**. Heat maps report the fold-change (Sz/control) for the 63 significantly altered synaptosome proteins in the synaptosome fraction (***Syn***), as well as the fold-change of those proteins in the homogenate fraction (***Hom***). Additionally, to assess the relationship between protein homogenate levels and synaptosome levels, we calculated synaptosome enrichment values (the ratio of a protein’s synaptosome levels over its homogenate levels) for each subject. ***Syn/Hom*** columns report the fold-change (Sz/control) for this metric. **C.** Lists the proteins for which the synaptosome enrichment values do not differ significantly between Sz and control; the synaptosome protein level differences in Sz are normalized by the homogenate level differences, suggesting that these synaptosome protein differences are driven by alterations in cell-wide protein expression (or degradation). **D.** Lists the proteins for which the synaptosome enrichment values differ between Sz and control significantly in the same direction as the synaptosome levels changes; the synaptosome level differences in Sz are not normalized by homogenate level changes, suggesting that mechanisms other than cell-wide protein expression (or degradation) are driving these effects.

### Effects of APD on protein levels

Of the 55 homogenate alterations, 10 showed alterations (uncorrected p < 0.05) in the same direction in APD monkey tissue (**Figure S6A, Table S7**). Of the 64 synaptosome alterations, one (ANK2) was similarly altered by APDs (p < 0.05) (**Figure S6B, Table S8**). Two proteins significantly decreased in Sz synaptosomes, GRIA2 and GBRB1, were increased (uncorrected p < 0.05) in haloperidol and/or olanzapine treated monkeys compared to vehicle (**Figure S6B**).

### Additional potential confounds

For a discussion of additional potential confounds in this cohort see *Supplement*.

### Protein co-regulation networks

The homogenate proteome organized into three interlinked modules (**Figure 4A & C**) enriched for terms relating to *Nucleus* and *Mitochondrion* (p = 7.5E^-6^, Turquoise), *Protein Transport* (p = 1.3E^-2^, Orange), and *Calmodulin-Binding* and *Cytoskeleton* (p = 2.1E^-4^ & 3E^-3^, Green). The *Nucleus* and *Mitochondrion* (Turquoise) module was enriched for proteins that differed between Sz and controls (p = 2.6E^-6^) and its Eigenprotein values differed significantly between Sz and controls (p = 2.6E^-4^, **Table S13**). **Figure 4E.** depicts the number of proteins in the Turquoise module that contribute to the selected gene ontologies, and the proportion of those that were altered in Sz, in a chord diagram.

**Figure 4.**
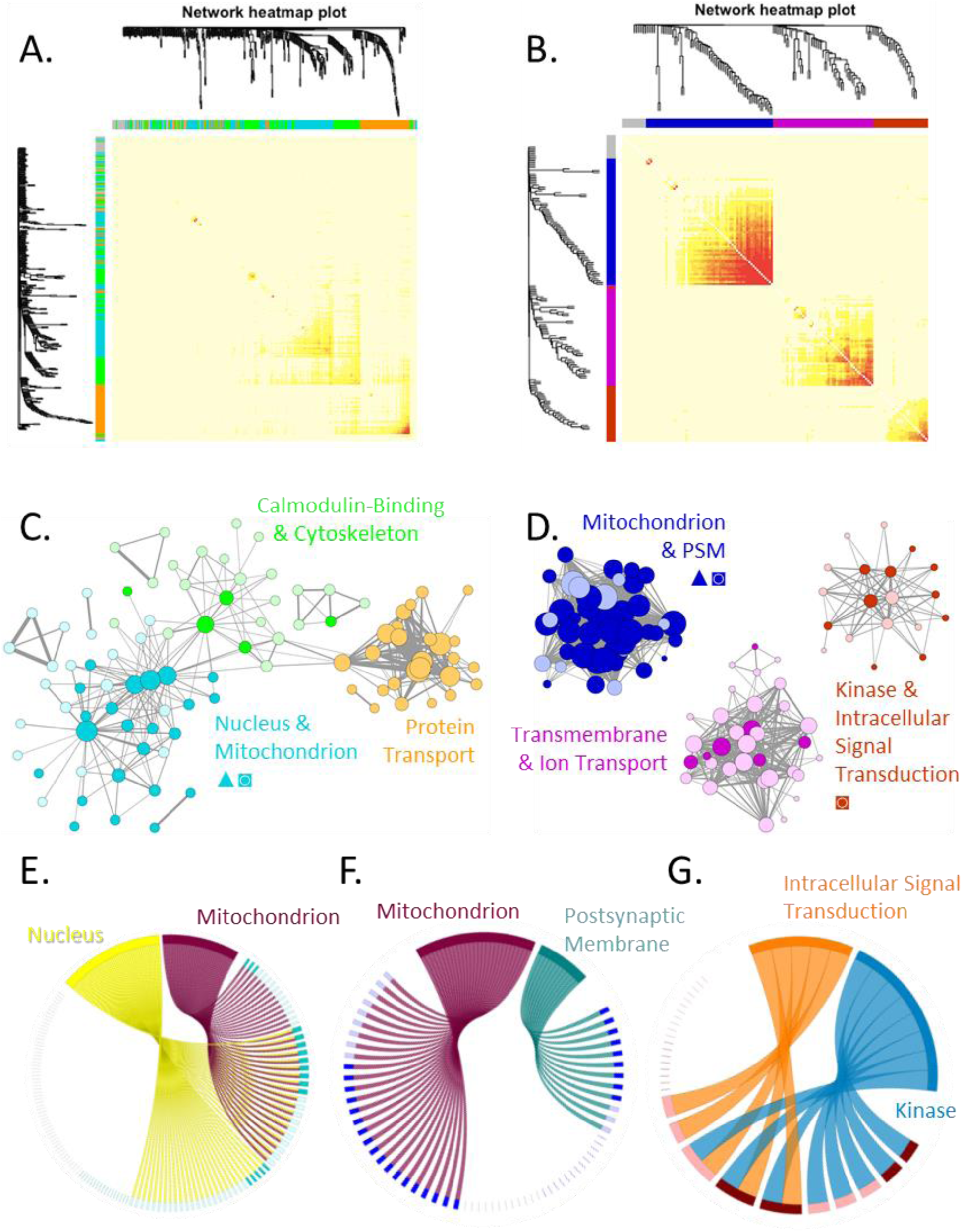
Homogenate and Synaptosome Protein Network Alterations. Protein co-regulation networks were constructed for the homogenate (**A.**) and synaptosome (**B.**) proteomes. Greater correlation between proteins is depicted in the heat maps as intensity of red color. Clustering of proteins into different modules is shown the by dendrograms and corresponding X and Y axis color coding. These networks were then visualized (**C.** homogenate, **D.** Control): Each node (circle) represents a protein. Color denotes module membership. Darker shading denotes proteins that were significantly altered in Sz. Node sizes are proportional to each proteins’ degree of connectivity. Line weights represent the strength of between-protein correlations. Only proteins with a weighted node connectivity > 0.15 were visualized. To investigate module biology, each module was first characterized by function/compartment using functional gene annotation analysis (e.g. Turquoise: *Nuclear* and *Mitochondrion*). Second, modules were tested for enrichment of Sz altered proteins, identifying the homogenate *Nuclear* and *Mitochondrion* module (Turquoise p = 6.2E^-6^) as well as the synaptosome *Mitochondrion & Postsynaptic Membrane* (PSM) (Blue p = 1.5E^-5^). Finally, we tested the difference in module Eigenprotein values between Sz and controls, identifying the Turquoise (7E^-5^), Blue (2.7E^-5^), and Red (0.015) modules as altered in Sz. ▴ = modules significantly enriched for proteins that differed significantly between Sz and control. ◙ = modules whose Eigenprotein values differed significantly between Sz and control. The chord figures depict the number of proteins in the Turquoise (E.), Blue (F.), and Red (G.) modules that contribute to the selected gene ontologies. Proteins contributing to ontologies are depicted as rectangles, with darker colors denoting those significantly altered in Sz. Proteins not contributing to the ontologies are depicted as lines, again with darker colors denoting significant alterations in Sz.

The synaptosome proteome organized into three highly distinct modules (**Figure 4B & D**) enriched for terms relating to *Mitochondrion* and *Postsynaptic Membrane* (p = 6.1E^-10^ & 2.9E^-3^, Blue), *Kinase* and *Intracellular Signal Transduction* (1.2E^-2^ & 4.6E^-2^, Red), and *Transmembrane* and *Ion Transport* (p = 1.4E^-8^ & 8.5E^-6^, Magenta). The *Mitochondrion* and *Postsynaptic Membrane* (Blue) module was enriched for proteins with levels that differed between Sz and control subjects (p = 1.5E^-5^) and its Eigenprotein values differed significantly between Sz and controls (p = 2E^-4^, **Table S13**). **Figure 4F**. depicts the numbers of proteins in the Blue module that contribute to the selected gene ontologies, as well as the proportion of those that were altered in Sz. In line with the analysis above, the 38 proteins whose synaptosome alterations were not explained by homogenate alterations (**Figure 3D**) were enriched for proteins from the *Mitochondrion* and *Postsynaptic Membrane* (Blue) module (p = 1.5E^-3^). Finally, Eigenprotein values for the *Kinase* and *Intracellular Signal Transduction* (Red) module differed between Sz and controls (p = 0.012, **Table S13**) (chord diagram: **Figure 4G**). Expanded chord diagrams for all homogenate and synaptosome modules, with protein identifiers and additional ontologies, are presented in **Figures S7-12.**

The Eigenprotein values from the *Calmodulin-Binding* and *Cytoskeleton* (Green) module were significantly inversely correlated with the *Protein Transport* (Orange) and *Mitochondrion* (Turquoise) modules (**Figure S13A**). Finally, we observed a significant correlation between the homogenate *Mitochondrion* (Turquoise) and synaptosome *Mitochondrion* and *Postsynaptic Membrane* (Blue) modules in the control, but not Sz, cohorts (**Figure S13B**).

### Similarities with reported transcriptome alterations

We did not observe a significant overlap between mRNA and homogenate protein alterations or mRNA and synaptosome protein alterations (**Figure S14A & B**). There was, however, a small but significant (r = 0.15, CI = 0.05048 to 0.2423, p = 3E^-3^) correlation between mRNA and homogenate protein Sz-control fold-changes (Figure S14C). No correlation was observed between mRNA and synaptosome protein Sz-control fold changes.

## Discussion

We observed robust differences in A1 homogenate and synaptosome protein levels in Sz. These differences organized into distinct co-regulation network alterations. Synaptosome levels of *Mitochondrion* and *Postsynaptic Membrane* proteins were generally lower in Sz, highly co-regulated, and did not appear to be driven by lower protein levels in the total homogenate. Homogenate levels of *Nuclear* and *Mitochondrion* proteins were lower in Sz, but were not correlated with synaptosome *Mitochondrion* and *Postsynaptic Membrane* proteins in disease. <u>Each</u> of these findings is discussed in turn below.

### Synaptosome Mitochondrion and Postsynaptic Membrane Proteins

Synaptosome alterations in the *Mitochondrion* and *Postsynaptic Membrane* module included a diverse array of proteins that were highly co-regulated, were mostly (but not uniformly) decreased in Sz, and are believed to localize to different synaptic microdomains. Specifically, this module included mitochondrial proteins (e.g. ADT3 and ATPA); ionotropic glutamate receptor subunits (NMDZ1, GRIA2, and GRIA3); GABA receptor subunits (GBRA1, GBRB1, and GBRG2); ion transporters (S12A5); adaptor proteins (ANK1 and ANK2); second messengers (RHG01); and a complement component (CO3). In the cortex, mitochondria are present in presynaptic boutons but not dendritic spines(36, 37); and ionotropic glutamate and GABA receptors are largely localized to different postsynaptic compartments. As we have not observed decreases in presynaptic boutons or inhibitory synapses in the A1 in Sz(38, 39), decreased synaptosome levels of mitochondrial proteins and GABA receptors are likely driven by alterations within their respective synaptic microdomains, not the loss of synaptic compartments. While we have observed decreased density of dendritic spines in Sz A1, we find it unlikely the observed decreases in synaptosome levels of postsynaptic glutamate receptors are driven by loss of the postsynaptic compartment as other canonical postsynaptic proteins (ANK1, ANK2, and RGH01) were increased and AMPA1 did not differ significantly. Placed in the context of what is known about Sz cytoarchitectonics, our findings suggest a robust, broad, and highly coordinated rearrangement of the synaptic proteome impacting both glutamate and GABA synapses as well as presynaptic bioenergetics. However, it is possible that a subset of our findings are driven by the loss of specific cellular or synaptic compartments in concert with altered protein levels in others. This question will ultimately have to be addressed by multiple-label confocal microscopy, RNA-scope, or electron microscopy studies, guided by our findings.

The majority of these synaptosome alterations were not explained by underlying changes in total homogenate protein levels, further suggesting that they are driven by alterations in local mitochondrial trafficking or synaptic proteostasis. Indeed, decreased mitochondrial number at axon terminals have been observed in Sz(40) and cellular mechanisms that regulate synaptic proteostasis are essential for synapse formation and plasticity and have previously been implicated in Sz. For example, the ubiquitin-proteasome system, a key regulator of synaptic protein stability, is essential for LTP and learning(41, 42) and has been implicated in Sz pathology by unbiased genetic analyses(43, 44). Thus, it is not surprising that ubiquitin-proteasome system alterations have been directly observed in Sz patient tissue(45-47). Similarly, phosphorylation is a well-documented regulator of glutamate receptor trafficking(48) and altered kinase activity has recently been observed in Sz(49, 50). In addition to posttranslational modification, alternative isoform expression and non-synonymous mutations can produce proteins with drastically different synaptic trafficking or anchoring properties(51, 52) and have also been recently implicated in Sz(53). In line with the results of these previous studies, our findings support a role for altered mitochondrial trafficking and synaptic proteostasis in Sz.

As neuronal excitation, inhibition, and bioenergetics are tightly balanced, it is not surprising that synaptic levels of these proteins are highly co-regulated. Considering the large body of work implicating postsynaptic glutamate receptor signaling in both Sz etiology and synaptic plasticity, it is a reasonable hypothesis that proteostatic alterations at excitatory postsynaptic compartments are the primary insult. For example, synaptosome levels of GRIA2, GRIA3, and S12A5 were decreased in Sz. Trafficking of GRIA2/3 containing AMPA receptors to new spines is essential for their potentiation and stabilization(54), as is synaptic S12A5(55-57). Subsequent decreases in synaptic mitochondrial and GABA receptors would then reflect homeostatic plasticity downstream of this decreased excitatory drive. Alternatively, presynaptic mitochondria are essential for bouton development(58) and activity (59). Thus, altered mitochondrial transport to the presynaptic compartment could also contribute to synaptic pathology in Sz.

### Cellular Homogenate Mitochondrion Proteins

We found reduced *Mitochondrion* protein levels in Sz cellular homogenates, consistent with numerous reports of lower tissue-wide expression of mitochondrial transcripts(44, 60-62), proteins(63), and enzymatic activity (64, 65) in the cortex of Sz subjects. As alterations in cell numbers have not been observed in Sz A1, these alterations likely reflect differences in cellular protein levels. Interestingly, cellular homogenate *Nuclear* and *Mitochondrion* Eigenprotein values and synaptosome *Mitochondrion* and *Postsynaptic Membrane* Eigenprotein values were highly correlated in controls, but not in Sz (**Figure S13**), indicating that cellular and synaptic mitochondria regulation are uncoupled in disease. This finding is consistent with the highly dynamic and responsive nature of the mitochondrial organelle; mitochondria localized to synaptic compartments would be specifically reactive to an altered synaptic micro-environment compared to mitochondria in other cellular domains.

### Relationship to transcriptome alterations

The homogenate protein alterations were weakly, but significantly, correlated with previously observed mRNA alterations in Sz(53), which is in line with multiple reports that mRNA and protein levels are only modestly correlated (66-68). Multiple processes beyond transcription regulate synaptic protein levels, such as RNA trafficking, translation, and protein trafficking. Thus, it is not surprising that synaptic protein alterations were not associated with previously reported mRNA alterations in Sz. This absence of a link between the synaptic proteome and the transcriptome (and, by extension, the genome) highlights a critical gap in our current understanding of Sz etiology and the need to investigate additional molecular features that regulate the synaptic proteome in patient tissue.

### Conclusion

We identified robust evidence of synaptic proteome alterations in the A1 of Sz. Some of these alterations appeared to be driven by tissue-wide changes, while others were exclusive to the synaptic compartment. Our findings are in-line with previous reports of postsynaptic glutamate receptor and mitochondrial impairments in Sz; identifying synapse specific alterations as well as providing for a link between the two. These findings provide a map to guide future studies into the upstream drivers of these synaptic impairments as well as their effects on specific cortical cells and circuits. Future proteomic studies will be required to determine how altered ubiquitination, phosphorylation, and isoform expression may each contribute to synaptic proteostasis impairments in Sz, while microscopy studies will be required to identify the cortical cell types and circuits to which these postsynaptic and mitochondrial alterations localize.

Dr. MacDonald had full access to all the data in the study and takes responsibility for the integrity of the data and the accuracy of the data analysis. Statistical analyses were performed by Dr. Ying Ding (University of Pittsburgh) and Dr. Matthew MacDonald (University of Pittsburgh).

## References

1. Javitt DC, Doneshka P, Grochowski S, Ritter W. Impaired mismatch negativity generation reflects widespread dysfunction of working memory in schizophrenia. Archives of General Psychiatry. 1995;52:550–558.

2. Javitt DC, Strous RD, Grochowski S, Ritter W, Cowan N. Impaired precision, but normal retention, of auditory sensory (“echoic”) memory information in schizophrenia. Journal of Abnormal Psychology. 1997;106:315–324.

3. Javitt DC, Shelley AM, Ritter W. Associated deficits in mismatch negativity generation and tone matching in schizophrenia. ClinNeurophysiol. 2000;111:1733–1737.

4. Rabinowicz EF, Silipo G, Goldman R, Javitt DC. Auditory sensory dysfunction in schizophrenia. Imprecision or distractibility? Archives of General Psychiatry. 2000;57:1149–1155.

5. Shelley AM, Silipo G, Javitt DC. Diminished responsiveness of ERPs in schizophrenic subjects to changes in auditory stimulation parameters: implications for theories of cortical dysfunction. Schizophrenia Research. 1999;37:65–79.

6. Ahveninen J, J„„skel„inen IP, Osipova D, Huttunen MO, Ilmoniemi RJ, Kaprio J, L”nnqvist J, Manninen M, Pakarinen S, Therman S, N„„t„nen R, Cannon TD. Inherited auditory-cortical dysfunction in twin pairs discordant for schizophrenia. Biological Psychiatry. 2006;60:612–620.

7. Hill SK, Beers SR, Kmiec JA, Keshavan MS, Sweeney JA. Impairment of verbal memory and learning in antipsychoticnaive patients with first-episode schizophrenia. Schizophr Res. 2004;68:127–136.

8. Mohn C, Torgalsboen AK. Details of attention and learning change in first-episode schizophrenia. Psychiatry Res. 2017;260:324–330.

9. Kantrowitz JT, Epstein ML, Beggel O, Rohrig S, Lehrfeld JM, Revheim N, Lehrfeld NP, Reep J, Parker E, Silipo G, Ahissar M, Javitt DC. Neurophysiological mechanisms of cortical plasticity impairments in schizophrenia and modulation by the NMDA receptor agonist D-serine. Brain. 2016;139:3281–3295.

10. Biagianti B, Fisher M, Neilands TB, Loewy R, Vinogradov S. Engagement with the auditory processing system during targeted auditory cognitive training mediates changes in cognitive outcomes in individuals with schizophrenia. Neuropsychology. 2016;30:998–1008.

11. Swerdlow NR, Bhakta SG, Light GA. Room to move: Plasticity in early auditory information processing and auditory learning in schizophrenia revealed by acute pharmacological challenge. Schizophr Res. 2018;199:285–291.

12. Moczulska KE, Tinter-Thiede J, Peter M, Ushakova L, Wernle T, Bathellier B, Rumpel S. Dynamics of dendritic spines in the mouse auditory cortex during memory formation and memory recall. Proc Natl Acad Sci U S A. 2013;110:18315–18320.

13. Yang Y, Liu DQ, Huang W, Deng J, Sun Y, Zuo Y, Poo MM. Selective synaptic remodeling of amygdalocortical connections associated with fear memory. Nat Neurosci. 2016;19:1348–1355.

14. Berry KP, Nedivi E. Spine Dynamics: Are They All the Same? Neuron. 2017;96:43–55.

15. Rosoklija G, Toomayan G, Ellis SP, Keilp J, Mann JJ, Latov N, Hays AP, Dwork AJ. Structural abnormalities of subicular dendrites in subjects with schizophrenia and mood disorders. Archives of General Psychiatry. 2000;57:349–356.

16. Garey LJ, Ong WY, Patel TS, Kanani M, Davis A, Mortimer AM, Barnes TR, Hirsch SR. Reduced dendritic spine density on cerebral cortical pyramidal neurons in schizophrenia. Journal of Neurology, Neurosurgery, and Psychiatry. 1998;65:446–453.

17. Glantz LA, Lewis DA. Decreased dendritic spine density on prefrontal cortical pyramidal neurons in schizophrenia. Archives of General Psychiatry. 2000;57:65–73.

18. Kolluri N, Sun Z, Sampson AR, Lewis DA. Lamina-specific reductions in dendritic spine density in the prefrontal cortex of subjects with schizophrenia. The American Journal of Psychiatry. 2005;162:1200–1202.

19. Sweet RA, Henteleff RA, Zhang W, Sampson AR, Lewis DA. Reduced dendritic spine density in auditory cortex of subjects with schizophrenia. Neuropsychopharmacology. 2009;34:374–389.

20. Konopaske GT, Lange N, Coyle JT, Benes FM. Prefrontal Cortical Dendritic Spine Pathology in Schizophrenia and Bipolar Disorder. JAMA psychiatry. 2014.

21. Schizophrenia Working Group of the Psychiatric Genomics C. Biological insights from 108 schizophrenia-associated genetic loci. Nature. 2014;511:421–427.

22. Alvarez-Castelao B, Schuman EM. The Regulation of Synaptic Protein Turnover. J Biol Chem. 2015;290:28623–28630.

23. Sutton MA, Schuman EM. Dendritic protein synthesis, synaptic plasticity, and memory. Cell. 2006;127:49–58.

24. Rosenberg T, Gal-Ben-Ari S, Dieterich DC, Kreutz MR, Ziv NE, Gundelfinger ED, Rosenblum K. The roles of protein expression in synaptic plasticity and memory consolidation. Front Mol Neurosci. 2014;7:86.

25. Lopez-Salon M, Alonso M, Vianna MR, Viola H, Melloe Souza T, Izquierdo I, Pasquini JM, Medina JH. The ubiquitin-proteasome cascade is required for mammalian long-term memory formation. Eur J Neurosci. 2001;14:1820–1826.

26. Lee SH, Choi JH, Lee N, Lee HR, Kim JI, Yu NK, Choi SL, Lee SH, Kim H, Kaang BK. Synaptic protein degradation underlies destabilization of retrieved fear memory. Science. 2008;319:1253–1256.

27. MacDonald ML, Ciccimaro E, Prakash A, Banerjee A, Seeholzer SH, Blair IA, Hahn CG. Biochemical fractionation and stable isotope dilution liquid chromatography-mass spectrometry for targeted and microdomain-specific protein quantification in human postmortem brain tissue. Mol Cell Proteomics. 2012;11:1670–1681.

28. Deo AJ, Cahill ME, Li S, Goldszer I, Henteleff R, Vanleeuwen JE, Rafalovich I, Gao R, Stachowski EK, Sampson AR, Lewis DA, Penzes P, Sweet RA. Increased expression of Kalirin-9 in the auditory cortex of schizophrenia subjects: Its role in dendritic pathology. Neurobiology of disease. 2011.

29. Deo AJ, Goldszer IM, Li S, DiBitetto JV, Henteleff RA, Sampson AR, Lewis DA, Penzes P, Sweet RA. PAK1 Protein Expression in the Auditory Cortex of Schizophrenia Subjects. PLoS One. 2013;8:e59458.

30. Konopaske GT, Dorph-Petersen KA, Sweet RA, Pierri JN, Zhang W, Sampson AR, Lewis DA. Effect of chronic antipsychotic exposure on astrocyte and oligodendrocyte numbers in macaque monkeys. Biol Psychiatry. 2008;63:759–765.

31. Chang-Gyu H, Anamika B, Mathew LM, Dan-Sung C, Joshua K, Zhiping N, Karin EB-W, Tilo G, Angel P, Eugene C, Steven EA, Hoau-Yan W, Blair IA. The Post-Synaptic Density of Human Postmortem Brain Tissues: An Experimental Study Paradigm for Neuropsychiatric Illnesses. Public Library of Sciences. 2009.

32. Wisniewski JR, Zougman A, Nagaraj N, Mann M. Universal sample preparation method for proteome analysis. Nat Methods. 2009;6:359–362.

33. MacLean B, Tomazela DM, Shulman N, Chambers M, Finney GL, Frewen B, Kern R, Tabb DL, Liebler DC, MacCoss MJ. Skyline: an open source document editor for creating and analyzing targeted proteomics experiments. Bioinformatics. 2010;26:966–968.

34. Law CW, Chen Y, Shi W, Smyth GK. voom: Precision weights unlock linear model analysis tools for RNA-seq read counts. Genome Biol. 2014;15:R29.

35. Langfelder P, Horvath S. WGCNA: an R package for weighted correlation network analysis. BMC Bioinformatics. 2008;9:559.

36. Santuy A, Turegano-Lopez M, Rodriguez JR, Alonso-Nanclares L, DeFelipe J, Merchan-Perez A. A Quantitative Study on the Distribution of Mitochondria in the Neuropil of the Juvenile Rat Somatosensory Cortex. Cereb Cortex. 2018;28:3673–3684.

37. AL. P. The fine structure of the nervous system: neurons and their supporting cells. Oxford University Press. 1991.

38. Moyer CE, Delevich KM, Fish KN, Asafu-Adjei JK, Sampson AR, Dorph-Petersen KA, Lewis DA, Sweet RA. Intracortical excitatory and thalamocortical boutons are intact in primary auditory cortex in schizophrenia SchizophrRes. 2013.

39. Moyer CE, Delevich KM, Fish KN, Asafu-Adjei JK, Sampson AR, Dorph-Petersen KA, Lewis DA, Sweet RA. Reduced Glutamate Decarboxylase 65 Protein Within Primary Auditory Cortex Inhibitory Boutons in Schizophrenia. Biol Psychiatry. 2012.

40. Roberts RC, Barksdale KA, Roche JK, Lahti AC. Decreased synaptic and mitochondrial density in the postmortem anterior cingulate cortex in schizophrenia. Schizophr Res. 2015;168:543–553.

41. Hegde AN. Proteolysis, synaptic plasticity and memory. Neurobiol Learn Mem. 2017;138:98–110.

42. Mabb AM, Ehlers MD. Ubiquitination in postsynaptic function and plasticity. Annu Rev Cell Dev Biol. 2010;26:179–210.

43. Liu C, Bousman CA, Pantelis C, Skafidas E, Zhang D, Yue W, Everall IP. Pathway-wide association study identifies five shared pathways associated with schizophrenia in three ancestral distinct populations. Transl Psychiatry. 2017;7:e1037.

44. Arion D, Corradi JP, Tang S, Datta D, Boothe F, He A, Cacace AM, Zaczek R, Albright CF, Tseng G, Lewis DA. Distinctive transcriptome alterations of prefrontal pyramidal neurons in schizophrenia and schizoaffective disorder. Mol Psychiatry. 2015;20:1397–1405.

45. Scott MR, Meador-Woodruff JH. Intracellular compartment-specific proteasome dysfunction in postmortem cortex in schizophrenia subjects. Mol Psychiatry. 2019.

46. Rubio MD, Wood K, Haroutunian V, Meador-Woodruff JH. Dysfunction of the ubiquitin proteasome and ubiquitinlike systems in schizophrenia. Neuropsychopharmacology. 2013;38:1910–1920.

47. Kim P, Scott MR, Meador-Woodruff JH. Abnormal expression of ER quality control and ER associated degradation proteins in the dorsolateral prefrontal cortex in schizophrenia. Schizophr Res. 2018.

48. Chen BS, Roche KW. Regulation of NMDA receptors by phosphorylation. Neuropharmacology. 2007;53:362–368.

49. McGuire JL, Depasquale EA, Funk AJ, O’Donnovan SM, Hasselfeld K, Marwaha S, Hammond JH, Hartounian V, Meador-Woodruff JH, Meller J, McCullumsmith RE. Abnormalities of signal transduction networks in chronic schizophrenia. NPJ Schizophr. 2017;3:30.

50. McGuire JL, Hammond JH, Yates SD, Chen D, Haroutunian V, Meador-Woodruff JH, McCullumsmith RE. Altered serine/threonine kinase activity in schizophrenia. Brain Res. 2014;1568:42–54.

51. Yang X, Coulombe-Huntington J, Kang S, Sheynkman GM, Hao T, Richardson A, Sun S, Yang F, Shen YA, Murray RR, Spirohn K, Begg BE, Duran-Frigola M, MacWilliams A, Pevzner SJ, Zhong Q, Trigg SA, Tam S, Ghamsari L, Sahni N, Yi S, Rodriguez MD, Balcha D, Tan G, Costanzo M, Andrews B, Boone C, Zhou XJ, Salehi-Ashtiani K, Charloteaux B, Chen AA, Calderwood MA, Aloy P, Roth FP, Hill DE, Iakoucheva LM, Xia Y, Vidal M. Widespread Expansion of Protein Interaction Capabilities by Alternative Splicing. Cell. 2016;164:805–817.

52. Richardson DS, Rodrigues DM, Hyndman BD, Crupi MJ, Nicolescu AC, Mulligan LM. Alternative splicing results in RET isoforms with distinct trafficking properties. Mol Biol Cell. 2012;23:3838–3850.

53. Gandal MJ, Zhang P, Hadjimichael E, Walker RL, Chen C, Liu S, Won H, van Bakel H, Varghese M, Wang Y, Shieh AW, Haney J, Parhami S, Belmont J, Kim M, Moran Losada P, Khan Z, Mleczko J, Xia Y, Dai R, Wang D, Yang YT, Xu M, Fish K, Hof PR, Warrell J, Fitzgerald D, White K, Jaffe AE, Psych EC, Peters MA, Gerstein M, Liu C, Iakoucheva LM, Pinto D, Geschwind DH. Transcriptome-wide isoform-level dysregulation in ASD, schizophrenia, and bipolar disorder. Science. 2018;362.

54. Man HY. GluA2-lacking, calcium-permeable AMPA receptors--inducers of plasticity? Curr Opin Neurobiol. 2011;21:291–298.

55. Llano O, Smirnov S, Soni S, Golubtsov A, Guillemin I, Hotulainen P, Medina I, Nothwang HG, Rivera C, Ludwig A. KCC2 regulates actin dynamics in dendritic spines via interaction with beta-PIX. J Cell Biol. 2015;209:671–686.

56. Li H, Khirug S, Cai C, Ludwig A, Blaesse P, Kolikova J, Afzalov R, Coleman SK, Lauri S, Airaksinen MS, Keinanen K, Khiroug L, Saarma M, Kaila K, Rivera C. KCC2 interacts with the dendritic cytoskeleton to promote spine development. Neuron. 2007;56:1019–1033.

57. Tornberg J, Voikar V, Savilahti H, Rauvala H, Airaksinen MS. Behavioural phenotypes of hypomorphic KCC2-deficient mice. Eur J Neurosci. 2005;21:1327–1337.

58. Lee CW, Peng HB. The function of mitochondria in presynaptic development at the neuromuscular junction. Mol Biol Cell. 2008;19:150–158.

59. Flippo KH, Strack S. An emerging role for mitochondrial dynamics in schizophrenia. Schizophr Res. 2017;187:26–32.

60. Iwamoto K, Bundo M, Kato T. Altered expression of mitochondria-related genes in postmortem brains of patients with bipolar disorder or schizophrenia, as revealed by large-scale DNA microarray analysis. Hum Mol Genet. 2005;14:241–253.

61. Vawter MP, Tomita H, Meng F, Bolstad B, Li J, Evans S, Choudary P, Atz M, Shao L, Neal C, Walsh DM, Burmeister M, Speed T, Myers R, Jones EG, Watson SJ, Akil H, Bunney WE. Mitochondrial-related gene expression changes are sensitive to agonal-pH state: implications for brain disorders. Mol Psychiatry. 2006;11:615, 663-679.

62. Middleton FA, Mirnics K, Pierri JN, Lewis DA, Levitt P. Gene expression profiling reveals alterations of specific metabolic pathways in schizophrenia. J Neurosci. 2002;22:2718–2729.

63. Karry R, Klein E, Ben Shachar D. Mitochondrial complex I subunits expression is altered in schizophrenia: a postmortem study. Biol Psychiatry. 2004;55:676–684.

64. Cavelier L, Jazin EE, Eriksson I, Prince J, Bave U, Oreland L, Gyllensten U. Decreased cytochrome-c oxidase activity and lack of age-related accumulation of mitochondrial DNA deletions in the brains of schizophrenics. Genomics. 1995;29:217–224.

65. Bergman O, Ben-Shachar D. Mitochondrial Oxidative Phosphorylation System (OXPHOS) Deficits in Schizophrenia: Possible Interactions with Cellular Processes. Can J Psychiatry. 2016;61:457–469.

66. de Sousa Abreu R, Penalva LO, Marcotte EM, Vogel C. Global signatures of protein and mRNA expression levels. Molecular bioSystems. 2009;5:1512–1526.

67. Zhang B, Gaiteri C, Bodea LG, Wang Z, McElwee J, Podtelezhnikov AA, Zhang C, Xie T, Tran L, Dobrin R, Fluder E, Clurman B, Melquist S, Narayanan M, Suver C, Shah H, Mahajan M, Gillis T, Mysore J, MacDonald ME, Lamb JR, Bennett DA, Molony C, Stone DJ, Gudnason V, Myers AJ, Schadt EE, Neumann H, Zhu J, Emilsson V. Integrated systems approach identifies genetic nodes and networks in late-onset Alzheimer’s disease. Cell. 2013;153:707–720.

68. Sharma K, Schmitt S, Bergner CG, Tyanova S, Kannaiyan N, Manrique-Hoyos N, Kongi K, Cantuti L, Hanisch UK, Philips MA, Rossner MJ, Mann M, Simons M. Cell type- and brain region-resolved mouse brain proteome. Nat Neurosci. 2015;18:1819–1831.

