## Supplement for "Synaptic Proteome Alterations in the Primary Auditory Cortex of Schizophrenia"

Supplement Contents:

1. Supplemental Tables S1-S18: legends and hyperlinks. (Pages 1 &2)
2. Supplemental Methods and Discussions (Pages 3-6)
3. Supplemental Figures S1-S14 with Legends (Pages 7- 22)

#### 1. Supplemental Tables:

**Table S1 Subject Demographics.** Reports demographic information for individual subjects.

**Table S2 Human Homogenate Protein Levels.** Reports the log2 protein levels for each subject, as well as group averages, and summary statistics for paired and limma voom analyses. Module membership for each protein is also reported.

**Table S3 Human Synaptosome Protein Levels.** Reports the log2 protein levels for each subject, as well as group averages, and summary statistics for paired and limma voom analyses. Module membership for each protein is also reported.

**Table S4 Synaptosome Enrichment.** To assess the enrichment (or de-enrichment) of each protein in the synaptosome preparations, the synaptosome level of each protein was divided by its homogenate level, for each subject in the control group. This table reports the averages of those enrichment values for each protein.

**Table S5 Mouse Homogenate PMI.** Reports the log2 homogenate protein levels for each sample as well as significance of the association between PMI and homogenate protein levels.

**Table S6 Mouse Synaptosome PMI.** Reports the log 2 synaptosome protein levels for each sample as well as significance of the association between PMI and homogenate protein levels.

**Table S7 Monkey APD Homogenate.** Reports the summary statistics for differences between the vehicle and haloperidol and vehicle and olanzapine groups.

**Table S8 Monkey APD Synaptosome.** Reports the log 2 synaptosome protein levels for each animal as well as summary statistics for differences between the vehicle and haloperidol and vehicle and olanzapine groups.

**Table S9 Eigenprotein Human Homogenate.** Reports the Eigenprotein values for the homogenate modules.

**Table S10 Eigenprotein Human Synaptosome.** Reports the Eigenprotein values for the synaptosome modules.

**Table S11 Homogenate Modules DAVID.** Reports the term enrichment for the homogenate modules.

**Table S12 Synaptosome Modules DAVID.** Reports the term enrichment for the synaptosome modules.

**Table S13 Module Eiegenprotein Differences.** Reports summary statistics for Sz-control differences in module Eigenproten values.

**Table S14 Aligned Sz-control Homogenate Proteome and Transcriptome Findings.** Reports the aligned adjusted p-values and Sz-control fold-changes from the homogenate proteome experiment in the present study and adjusted p-values and Sz-control fold-changes from a meta-analysis of transcriptome studies in Sz tissue (Gandel et al. 2018).

**Table S15 Aligned Sz-control Homogenate Proteome and Transcriptome Findings.** Reports the aligned adjusted p-values and Sz-control fold-changes from the synaptosome proteome experiment in the present study and adjusted p-values and Sz-control fold-changes from a meta-analysis of transcriptome studies in Sz tissue (Gandel et al. 2018).

**Table S16 Association of Potential Confounds and Homogenate Protein Levels.** Reports associations between potential confounds (pH, PMI, Age, Storage Time, Tobacco, Other medications, Suicide, Schizoaffective diagnosis, Benzodiazepines, Anticonvulsants, Antidepressants, Antipsychotic, Alcohol of substance abuse, Duration, Age of onset, Sex, and Cannabis) and homogenate protein levels.

**Table S17 Association of Potential Confounds and Synaptosome Protein Levels.** Reports associations between potential confounds (pH, PMI, Age, Storage Time, Tobacco, Other medications, Suicide, Schizoaffective diagnosis, Benzodiazepines, Anticonvulsants, Antidepressants, Antipsychotic, Alcohol of substance abuse, Duration, Age of onset, Sex, and Cannabis) and synaptosome protein levels.

**Table S18 List of Protein SRMs from Assay Stages.** Reports the number of proteins for which SRMs were 1. Assayed in the MS run; 2. Passed quality control; 3. Were altered between Sz and control (unadjusted Limma $p < 0.05$); and 4. Were altered between Sz and control (adjusted Limma $p < 0.05$).

#### 2. Supplemental Methods and Discussions:

**Human Subjects and Tissue Collection:** Brain specimens from all subjects were obtained during autopsies conducted at the Allegheny County Office of the Medical Examiner after receiving consent from the next-of-kin. An independent panel of experienced clinicians made consensus Diagnostic and Statistical Manual of Mental Disorders Fourth Edition (DSM-IV) diagnoses using a previously described method(1). 48 pairs of Sz and control subjects (matched by age, sex, race, and PMI, **Table 1 & S1**) were studied. Control subjects underwent identical assessments and were determined to be free of lifetime psychiatric illness. Procedures were approved by the University of Pittsburgh Institutional Review Board and Committee for Oversight of Research Involving the Dead. Grey matter was harvested from the auditory cortex as previously described(2, 3): Tissue slabs containing the superior temporal gyrus with Heschl's Gyrus (HG) located medial to the planum temporal were identified, and the superior temporal gyrus (STG) removed as single block. The samples were then distributed in a block design for preparation and analysis to evenly distribute SCZ and control samples and blind the experimenters during sample preparation, analysis, and peak integration. Grey matter was collected from HG by taking 40 μm sections and frozen at -80 °C(2).

**Antipsychotic Drug Treated Monkey Tissue:** The tissue utilized here was obtained from previously completed experiments that have been extensively described previously(4). Briefly: 18 male monkeys, n = 6 per group, were administered therapeutically relevant doses of haloperidol, olanzapine, or vehicle. Tissue slabs containing the superior temporal gyrus were removed in a single block and 50 mg of grey matter was harvested as described above.

**Sample Preparation:** Total Homogenate and Synaptosome preparations were obtained using a variation on the sucrose density gradient centrifugation method we have previously validated for use in human postmortem brain tissue(5, 6). Briefly: 50 mg of human A1 grey matter, 50 mg monkey STG grey matter, or one mouse brain hemisphere were homogenized in 0.32 M Sucrose, 0.1 mM CaCl2, 1mM MgCl2 with 1 x Protease and Phosphate Inhibitors (Sigma) (10 µl/mg brain tissue) on ice in a 1 ml glass Dounce homogenizer with a Teflon pestle. 40 µl of this crude homogenate is mixed with 10 µl 10% SDS, vortexed, sonicated, and cleared by centrifugation to prepare the “Total Homogenate” (Hom). The remainder of crude homogenate is clarified at 800 x g for 10 min at 4C and 750 µl of the supernatant is adjusted to 1.25 M sucrose in a final volume of 1875 µl. The 1.25 M sucrose-homogenate is laid between 5 ml of 1.5 M sucrose (bottom) and 1.5 ml of 1 M sucrose (top) and spun at 24,000 RPM for 3 hrs (4C) in a TH641 swing arm rotor. The band at the 1.25-1.5 M interface is collected into 10 ml 0.05 mM $CaCl_2$ and spun at 18,000 RPM for 20 min in a TH641 swing arm rotor. The pellet is taken up in 40 µl 100 mM Tris, pH 7.4, 2% SDS (1X Protease and Phosphate Inhibitors, Sigma), vortexed, sonicated, and cleared by centrifugation to prepare the “Synaptosomes” (Syn). Total protein concentration in the Hom and Syn preparations were assessed by micro BCA (Pierce).

**[$^{13}C_6$]brain ISTD:** The [$^{13}C_6$]lysine-labeled brain proteome internal standard ([$^{13}C_6$]brain ISTD) is prepared by homogenizing cerebral cortex tissue from a Stable Isotope Labeling in Mammals (SILAM) mouse (Cambridge Isotopes). These animals are raised on a diet in which the only source of Lysine is $^{13}C_6$ labeled, resulting in near complete (99%) labeling of the animal proteome in three generations. Labeling efficiency of each [$^{13}C_6$]brain ISTD preparation is confirmed prior to use. To account for species differences, SRMs are designed only for peptides that have 100% homology between humans and mice(5) taking advantage of the high level of synaptic protein sequence homology between these species (7).

**Peptide SRM Development and Validation:** Method development began by building upon our previously described SRM libraries(5, 8-10) with the selection of additional proteins of interest included in published multidimensional MS/MS analyses of synaptic enrichments from mouse and human brain tissue (5, 6, 11) as well as the NIST Tandem Mass Spectral Library. Targets for inclusion in the LC-SRM/MS assay were selected with a bias toward well annotated synaptic proteins, such as glutamate receptors, kinases, phosphatases, vesicular fusion, amino acid metabolism, protein trafficking and scaffolding as well as proteins implicated by recent unbiased genetic analyses of Sz(12). Peptides for proteins of interest were then filtered based on the following criteria: 1) presence of lysine, 2) nonredundant to a selected protein or protein group (determined by BLAST search) and 3) 100% homology across mouse and human sequences (determined by BLAST search). Ultimately, 2700 tryptic peptides from 152 proteins were selected for validation for addition to the assay. Acceptable peptide sequences, along with MS2 spectra, were imported into Skyline(13). Initially, five mass transitions were selected for each target peptide and its “heavy” counter-part. Validation experiments were performed in 1:1 (µg/ µg) mixtures of a human synaptosome preparation from a low PMI control and the [$^{13}C_6$]brain ISTD. To ensure that the desired peptide was assayed, rigorous selection criteria were employed for the inclusion of peptide SRM transitions. Candidate “light”/“heavy” peptide SRMs were evaluated manually in Skyline for A) retention time, B) similar y-ion ratios (within 25%) to each other and database MS2 spectra, and D) a signal-to-noise ratio greater than 3. Peptide SRM pairs for which more than one identical peak was observed were omitted. Ultimately, SRMs for 825 peptides unique to 478 proteins (combined from this round of method development and our published studies) were validated at time of assay.

**Reproducibility of Peptide Quantification and Calculation of Protein Level Measures:** To assess variabilities from sample preparation and MS analysis, coefficients of variation (CVs) were calculated for each data cohort based on peptide values from the enrichment, pooled controls, and MS controls (**Figure 1**). Peptides with CVs > 0.5 in the MS control were discarded. For multiple peptides mapped to the same protein, their mean abundance value, inversely weighted based on the sum of CVs from all three of the control series, was used as the protein-level measure for the downstream analysis. Of the peptide SRMs described above, 777 unique to 408 proteins were quantified in the human homogenates and 575 peptides unique to 351 proteins (a subset of the 408 homogenate proteins) were quantified in human synaptosome. Mean CV for peptide quantification in the human experiments was 0.12 (SD = 0.09) in the homogenates, 0.17 in synaptosomes (SD = 0.12) (**Figure S2**).

**Differential Expression Analysis**: Limma-Voom(14) was performed to detect the difference between Sz and control samples in homogenate and synaptosome proteins, respectively. We estimated the correlation between paired subject samples using R function “duplicateCorrelation”. Empirical Bayes statistics were used to rank the proteins in the order of evidence for differential expression. The moderated t-statistics (based on the moderated standard errors for increased statistical power), p-values and q-values were obtained (Tables S2 and S3). Similar analyses using Limma-Voom were performed with pH and pH + Age added as covariates. T-tests were performed on protein data from monkey samples to compare haloperidol or olanzapine group versus the vehicle group.

**Effects of PMI:** One-way ANOVA was used to assess the effects of PMI on homogenate and synaptosome protein levels, respectively. The levels of all proteins for which an effect of PMI was detected (uncorrected $p < 0.05$, **Tables S5 & S6**), were plotted versus PMI and inspected by the senior author, blinded to protein identification. Proteins that displayed a step-wise increase or decrease (**Figures S3 & S4**) were discarded while those that displayed linear or curved increase or decrease (and could presumably be controlled for by PMI matching) were retained (**Figures S3**). PMI had an effect (uncorrected $p < 0.05$) on the levels of 60 proteins in homogenate and 70 proteins in synaptosomes (**Tables S5 & S6**). Of these, six in homogenate (B4DV12, E41L3, GFAP, PRDX1, TPIS, and DYN3) and four in synaptosomes (HS12A, KPCA, SYN1, and 2A5D) displayed step-wise alterations across PMI (**Figures S3 & S4**) and were dropped from analysis.

**Network Analysis:** Weighted Gene Co-expression Network Analysis (WGCNA)(34) was used to investigate the pattern of co-regulated protein expression. First, for homogenate and enriched synaptosome proteins, we performed exploratory WGCNA on Sz and control samples, separately. Then, we conducted module preservation analyses to test whether the modules identified from the Sz samples were preserved in the control samples, or vice versa (Figure S5) (35). Since all the modules detected from Sz samples were modestly to strongly preserved in the control samples and vice versa, we constructed protein co-expression networks using combined Sz and control samples, for homogenate (402 proteins) and enriched synaptosome (155 enriched proteins), respectively. Module membership was characterized for Gene Ontology terms by functional enrichment using DAVID(36). The two networks were visualized using Cytoscape. To quantitate the representation of protein levels in each module, we calculated the first principal component of modules given by WGCNA, referred to as the module-specific Eigenprotein (**Tables S9 & S10**). Sz-control differences between module-specific Eigenproteins were assessed by Student's t-test. A linear regression model was fitted to model the correlation between the Eigenproteins of each module, with diagnosis and batch included as covariates.

**Comparison to PsychENCODE mRNA Findings:** Homogenate and Synaptosome protein findings from this study were compared to findings from a recent meta-analysis(15) of RNAseq studies from the PsychENCODE Consortium(16). This study integrated analyses of frontal and temporal cerebral cortex tissue from 559 Sz subjects and 936 controls. Ensembl identifiers from the RNAseq meta-analysis were mapped to UniProtKB identifiers in our data using the *Retrieve/ID mapping* tool(17), and reported fold-changes and adjusted p-values for Sz-control transcript differences extracted for the aligned proteins (**Tables S14 & S15**). Of the 402 proteins investigated in the present study, all but two (GCG: glucagon and SPTA1: Spectrin Alpha, Erythrocytic 1, proteins for which transcripts would not be expected to be present in brain tissue) were mapped mRNA transcripts. Fisher's exact test was used to assess if a significant number of homogenate or synaptosome protein alterations overlapped with the reported transcriptome differences (**Figure S14A & B**). Additionally, linear regressions were run to assess the level of association between mRNA-homogenate protein level fold-changes and mRNA-synaptosome protein level fold-changes (**Figures S14C & D**).

**Additional Potential Confounds:** In addition to antipsychotic medication and PMI, several other potential confounding factors were present in this cohort, including (but not limited to) pH, age, storage time, smoking tobacco, death by suicide, and schizoaffective disorder (**Tables S16 and S17**). Of these, only age significantly correlated with homogenate protein levels, and only age and pH significantly correlated with synaptosome protein levels ($q < 0.05$, Tables S16 and S17). Based on these findings, we ran addition Limma-voom case-control analyses including **1.** pH as a co-variate and **2**. pH and age as covariates. The results of both of these analyses are included in **Tables S2 and S3**. Age did not impact our findings. Inclusion of pH had little overall effect on the associations of protein levels with diagnosis in synaptosome. Although the inclusion of this covariate did slightly increase the smallest p-values and the corresponding adjusted p-values, the rank order of association in the two analyses were highly correlated (**Table S3**, columns L and P, **Figure S15**). We have chosen to present the Limma-voom without pH as our primary analysis as pH cannot be disentangled from disease process. As all the tissue included this study was obtained from individuals without prolonged agonal states (as reflected in the overall high pH across samples), pH differences are not a result of a technical or sampling artifact. Instead, mitochondrial alterations within individuals with Sz, evidenced by altered mitochondrial protein levels in both homogenate and synaptosome samples in our studies, would be expected to alter tissue pH.


1. Glantz LA, Lewis DA. Decreased dendritic spine density on prefrontal cortical pyramidal neurons in schizophrenia. Archives of General Psychiatry. 2000;57(1):65-73.
2. Deo AJ, Cahill ME, Li S, Goldszer I, Henteleff R, Vanleeuwen JE, et al. Increased expression of Kalirin-9 in the auditory cortex of schizophrenia subjects: Its role in dendritic pathology. Neurobiology of disease. 2011(Journal Article).
3. Deo AJ, Goldszer IM, Li S, DiBitetto JV, Henteleff RA, Sampson AR, et al. PAK1 Protein Expression in the Auditory Cortex of Schizophrenia Subjects. PLoS One. 2013;8(4):e59458.
4. Konopaske GT, Dorph-Petersen KA, Sweet RA, Pierri JN, Zhang W, Sampson AR, et al. Effect of chronic antipsychotic exposure on astrocyte and oligodendrocyte numbers in macaque monkeys. Biol Psychiatry. 2008;63(8):759-65.
5. MacDonald ML, Ciccimaro E, Prakash A, Banerjee A, Seeholzer SH, Blair IA, et al. Biochemical fractionation and stable isotope dilution liquid chromatography-mass spectrometry for targeted and microdomain-specific protein quantification in human postmortem brain tissue. Mol Cell Proteomics. 2012;11(12):1670-81.
6. Chang-Gyu H, Anamika B, Mathew LM, Dan-Sung C, Joshua K, Zhiping N, et al. The Post-Synaptic Density of Human Postmortem Brain Tissues: An Experimental Study Paradigm for Neuropsychiatric Illnesses. Public Library of Sciences. 2009(Journal Article).
7. Bayes A, van de Lagemaat LN, Collins MO, Croning MD, Whittle IR, Choudhary JS, et al. Characterization of the proteome, diseases and evolution of the human postsynaptic density. Nature Neuroscience. 2011;14(1):19-21.
8. MacDonald ML, Favo D, Garver M, Sun Z, Arion D, Ding Y, et al. Laser capture microdissection-targeted mass spectrometry: a method for multiplexed protein quantification within individual layers of the cerebral cortex. Neuropsychopharmacology. 2019;44(4):743-8.

9. Krivinko JM, Erickson SL, Ding Y, Sun Z, Penzes P, MacDonald ML, et al. Synaptic Proteome Compensation and Resilience to Psychosis in Alzheimer's Disease. Am J Psychiatry. 2018;175(10):999-1009.
10. MacDonald ML, Ding Y, Newman J, Hemby S, Penzes P, Lewis DA, et al. Altered glutamate protein co-expression network topology linked to spine loss in the auditory cortex of schizophrenia. Biol Psychiatry. 2015;77(11):959-68.
11. Trinidad JC, Thalhammer A, Specht CG, Lynn AJ, Baker PR, Schoepfer R, et al. Quantitative analysis of synaptic phosphorylation and protein expression. Molecular {\&} Cellular Proteomics. 2008;7(4):684-96.
12. Lachman HM, Morrow B, Shprintzen R, Veit S, Parsia SS, Faedda G, et al. Association of codon 108/158 catechol-O-methyltransferase gene polymorphism with the psychiatric manifestations of velo-cardio-facial syndrome. American Journal of Medical Genetics. 1996;67(5):468-72.
13. MacLean B, Tomazela DM, Shulman N, Chambers M, Finney GL, Frewen B, et al. Skyline: an open source document editor for creating and analyzing targeted proteomics experiments. Bioinformatics. 2010;26(7):966-8.
14. Law CW, Chen Y, Shi W, Smyth GK. voom: Precision weights unlock linear model analysis tools for RNA-seq read counts. Genome Biol. 2014;15(2):R29.
15. Gandal MJ, Zhang P, Hadjimichael E, Walker RL, Chen C, Liu S, et al. Transcriptome-wide isoform-level dysregulation in ASD, schizophrenia, and bipolar disorder. Science. 2018;362(6420).
16. Psych EC, Akbarian S, Liu C, Knowles JA, Vaccarino FM, Farnham PJ, et al. The PsychENCODE project. Nat Neurosci. 2015;18(12):1707-12.
17. UniProt C. UniProt: a worldwide hub of protein knowledge. Nucleic Acids Res. 2019;47(D1):D506-D15.

#### 3. Supplemental Figures:

**Figure S1. Synaptosome yield, PMI, and pH in human tissue.** Linear regression analyses of synaptosome yield (µg total synaptosome protein / mg tissue) with PMI (**A**) and pH (**B**).

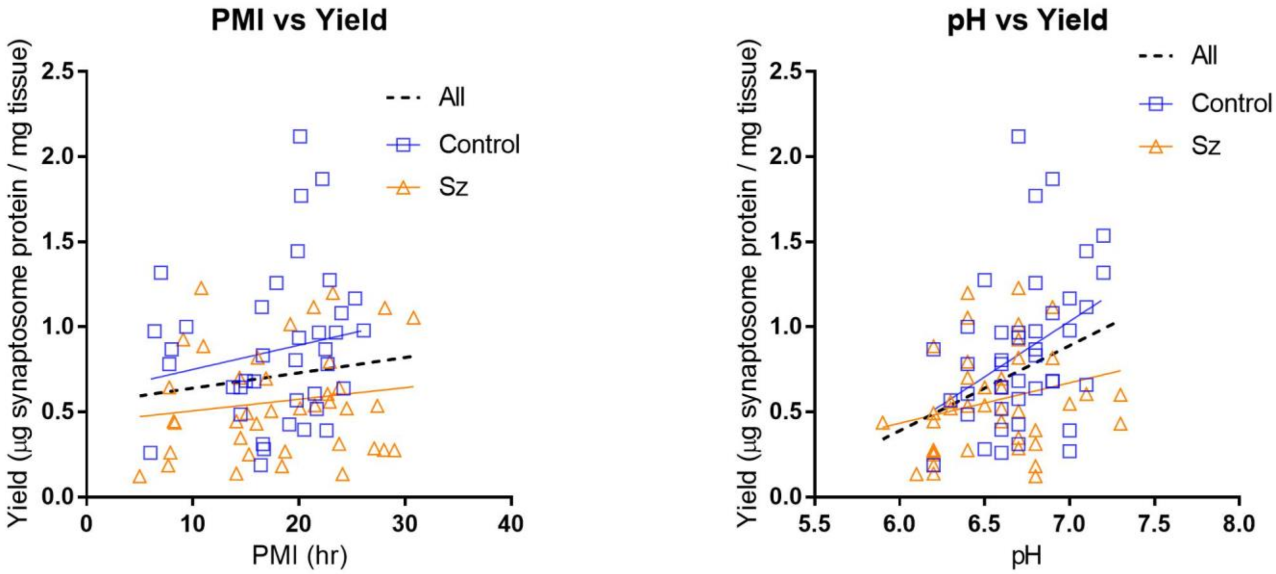

**Figure S2. Distribution of Technical and Variance in Peptide Quantification. A.** 777 peptides unique to 402 proteins were quantified in 12 homogenate Enrichment Controls. The mean CV was 0.12 with 75% peptide CVs < 0.17. **B.** 575 peptides unique to 348 proteins were quantified in 12 synaptosome Enrichment Controls. The mean CV was 0.17 with 75% peptide CVs < 0.19 **C.** 777 peptides unique to 402 proteins were quantified in 8 homogenate Trypsin Controls. The mean CV was 0.08 with 75% peptide CVs < 0.13. **D.** 575 peptides unique to 348 proteins were quantified in 8 synaptosome Trypsin Controls. The mean CV was 0.1 with 75% peptide CVs < 0.13. **E.** 777 peptides unique to 402 proteins were quantified in 8 homogenate MS Controls. The mean CV was 0.1 with 75% peptide CVs < 0.13.

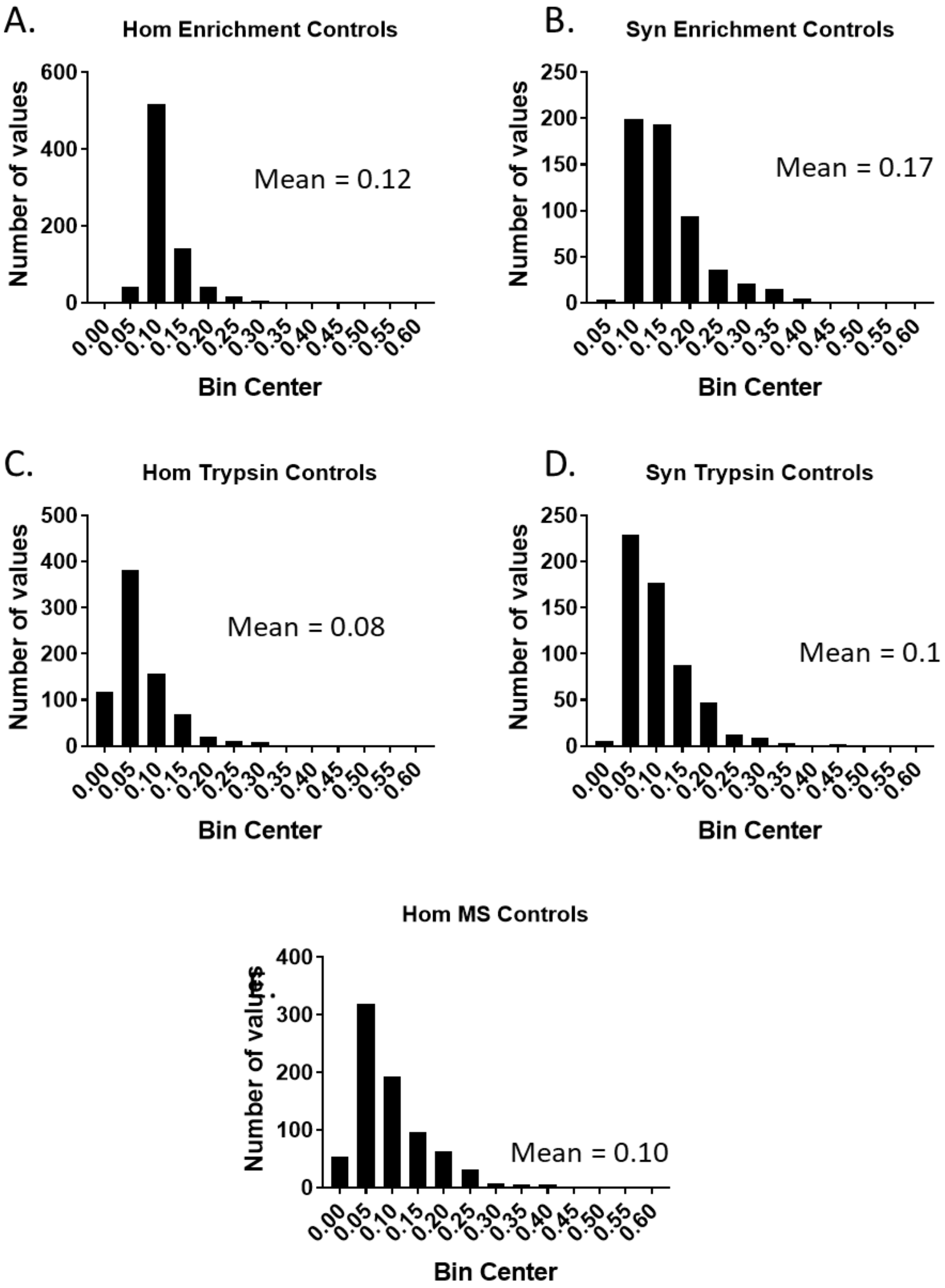

**Figure S3. Homogenate Protein Levels and PMI in Mice. A. – F.** show the six proteins with homogenate levels that were significantly correlated with PMI in mice with stepwise changes. These proteins were dropped from analysis. **G. - I.** show representative proteins that were retained in statistical analysis.

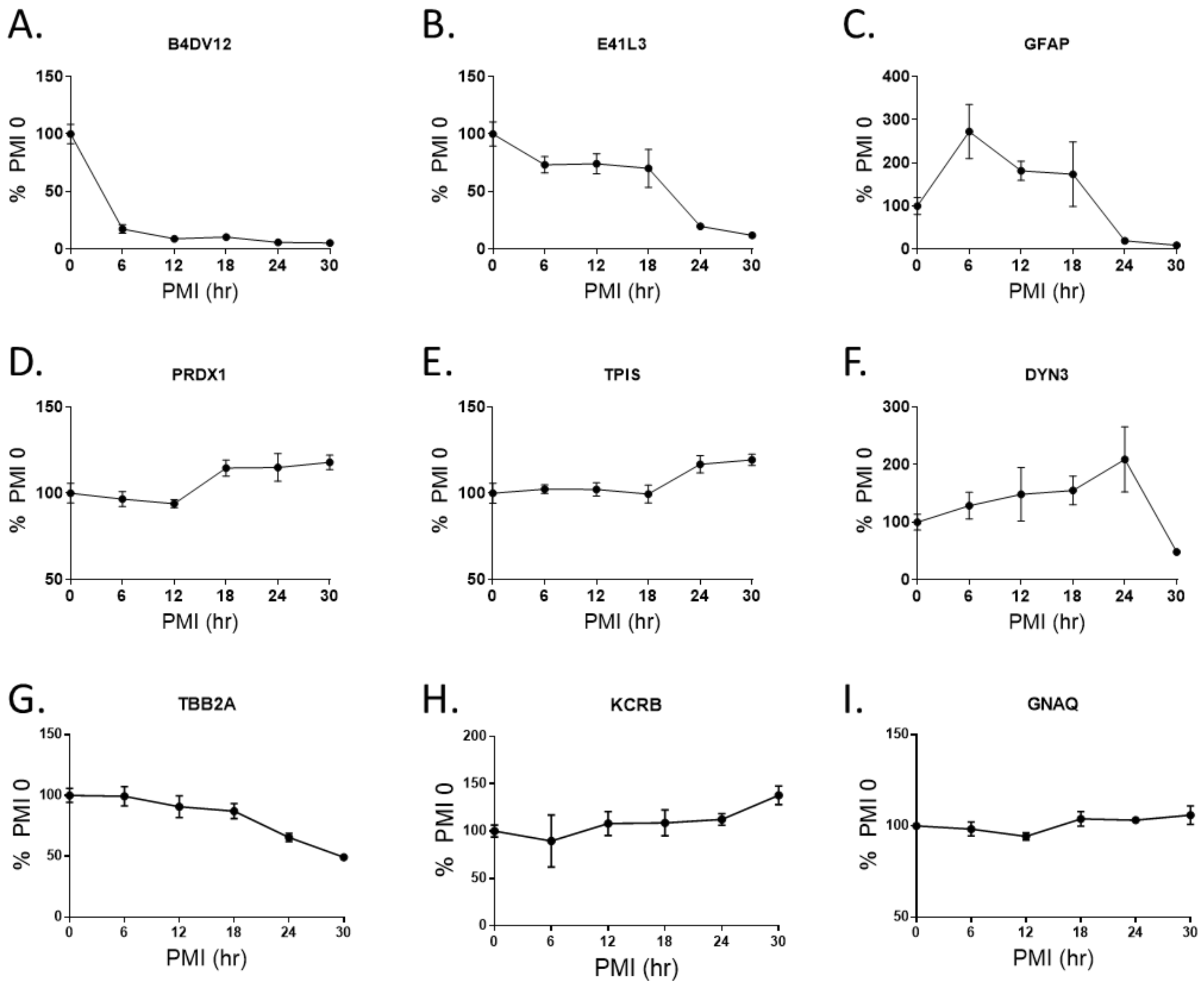

**Figure S4. Synaptosome Protein Levels and PMI A. – D.** show the six proteins with synaptosome levels that were significantly correlated with PMI in mice with stepwise changes. These proteins were dropped from analysis.

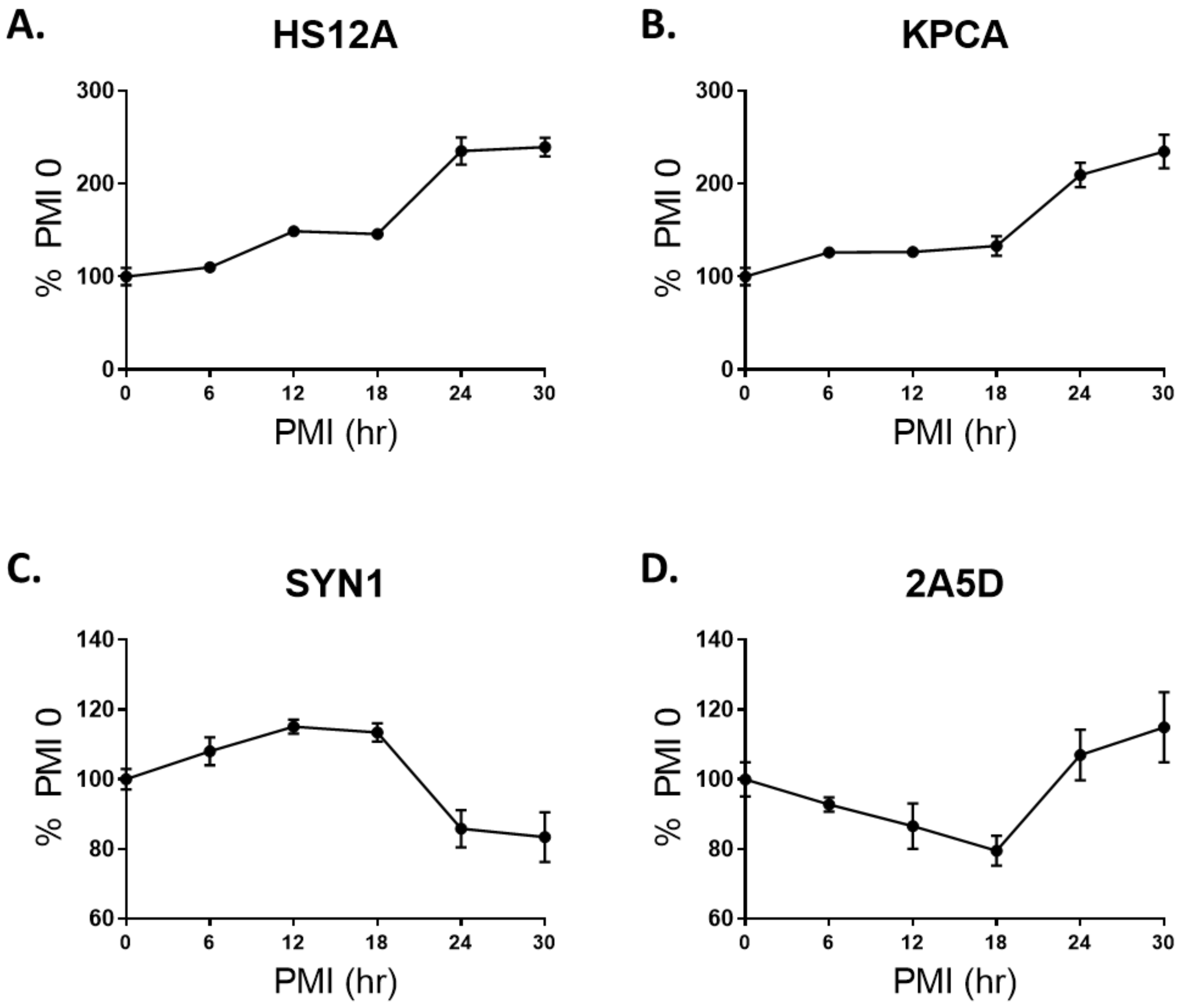

**Figure S5. Exploratory Module Preservation Analysis.** Panels **A.**, **C.**, **E.**, and **G.** show the composite statistic medianRank versus module size. The higher the medianRank, the less preserved is the module relative to other modules. Panels **B.**, **D.**, **F.**, and **H.** show the composite statistic Zsummary versus module size. Zsummary < 2 indicates no preservation, 2 < Zsummary < 10 indicates weak to moderate evidence of preservation, Zsummary > 10 indicates strong evidence of preservation. This exploratory analysis utilized different WGCNA parameters than subsequent WGCNA builds, allowing for a greater number of modules.

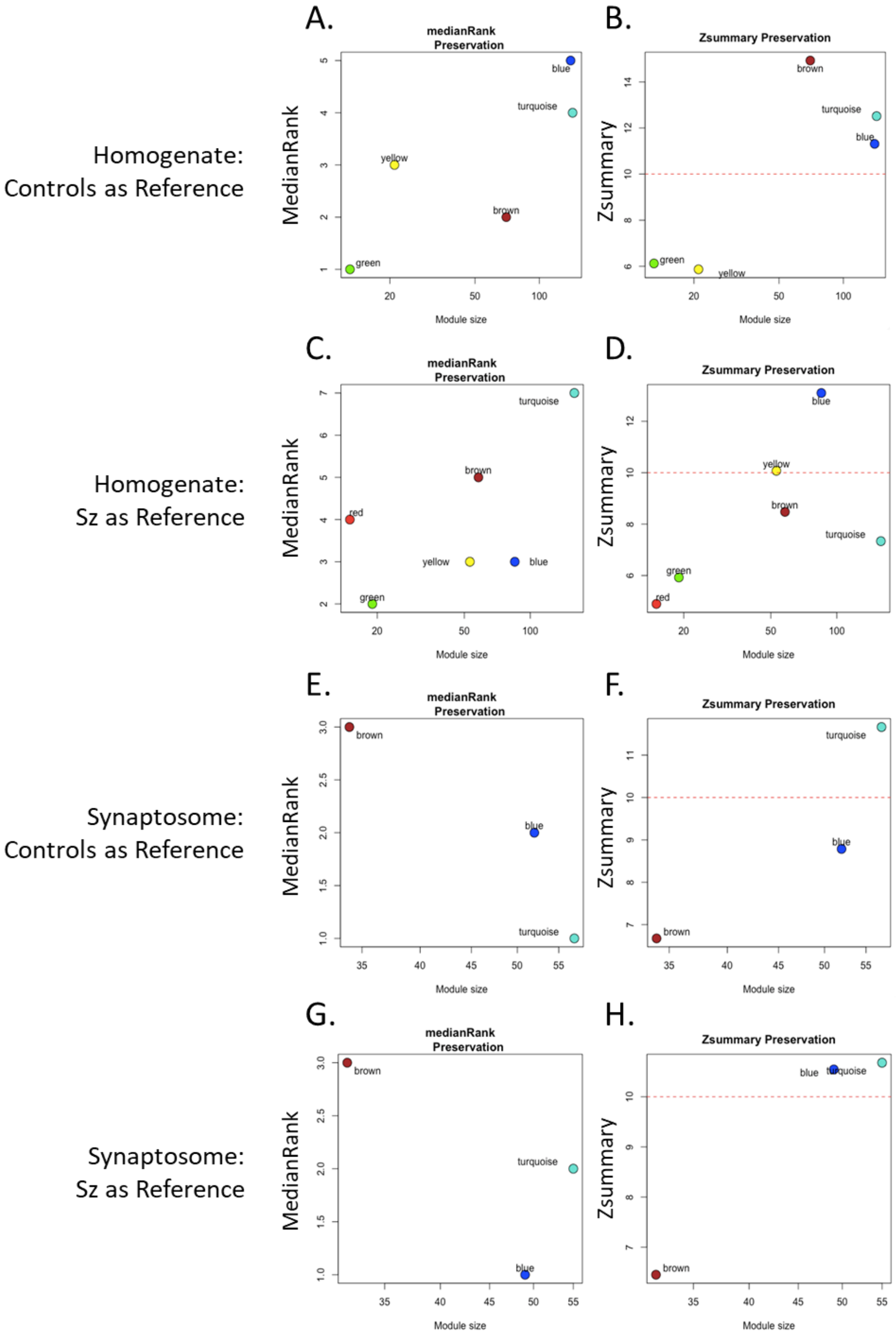

**Figure S6. Comparison of antipsychotic drug effects and Sz alterations.** These heat maps report fold-changes between Sz and control (Sz), haloperidol and vehicle (Hal), and olanzapine and vehicle (Olz) in homogenates (**A.**) and synaptosomes (**B.**). Windows with thicker boarders denote significant fold-changes in Sz (q < 0.05) and nominally significantly fold-changes in Hal and Olz (p < 0.05).

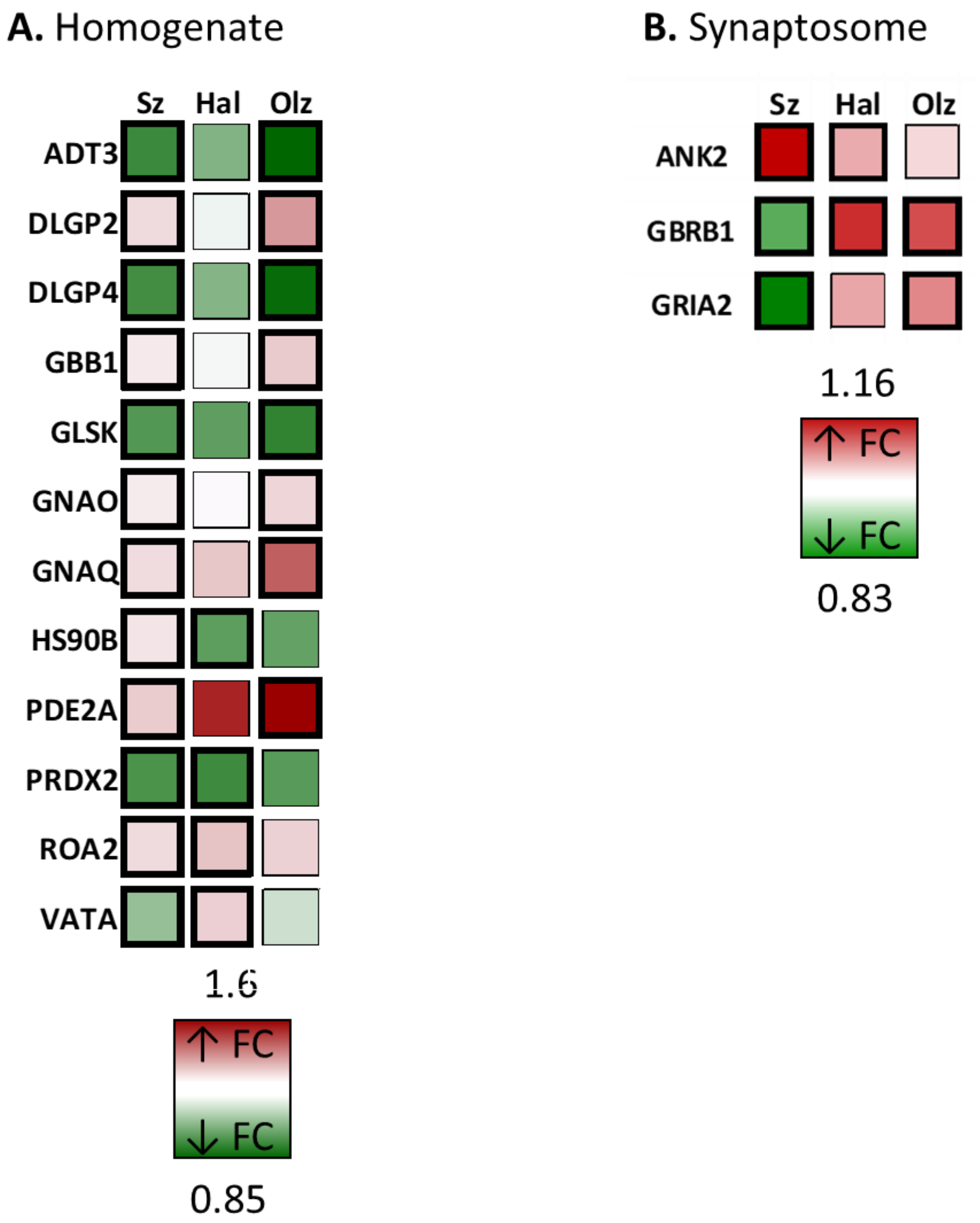

**Figures S7-S12. Expanded Chord Diagrams for All Homogenate and Synaptosome Modules.** The chord figures depict the number of proteins in each module that contribute to the selected gene ontologies. p-values report the significance of enrichment for the ontology term in the modules. Proteins contributing to ontologies are depicted as rectangles, with darker colors denoting those significantly altered in Sz. Proteins not contributing to the ontologies are depicted as lines, again with darker colors denoting significant alterations in Sz.

### Figure S7.
#### Homogenate Turquoise

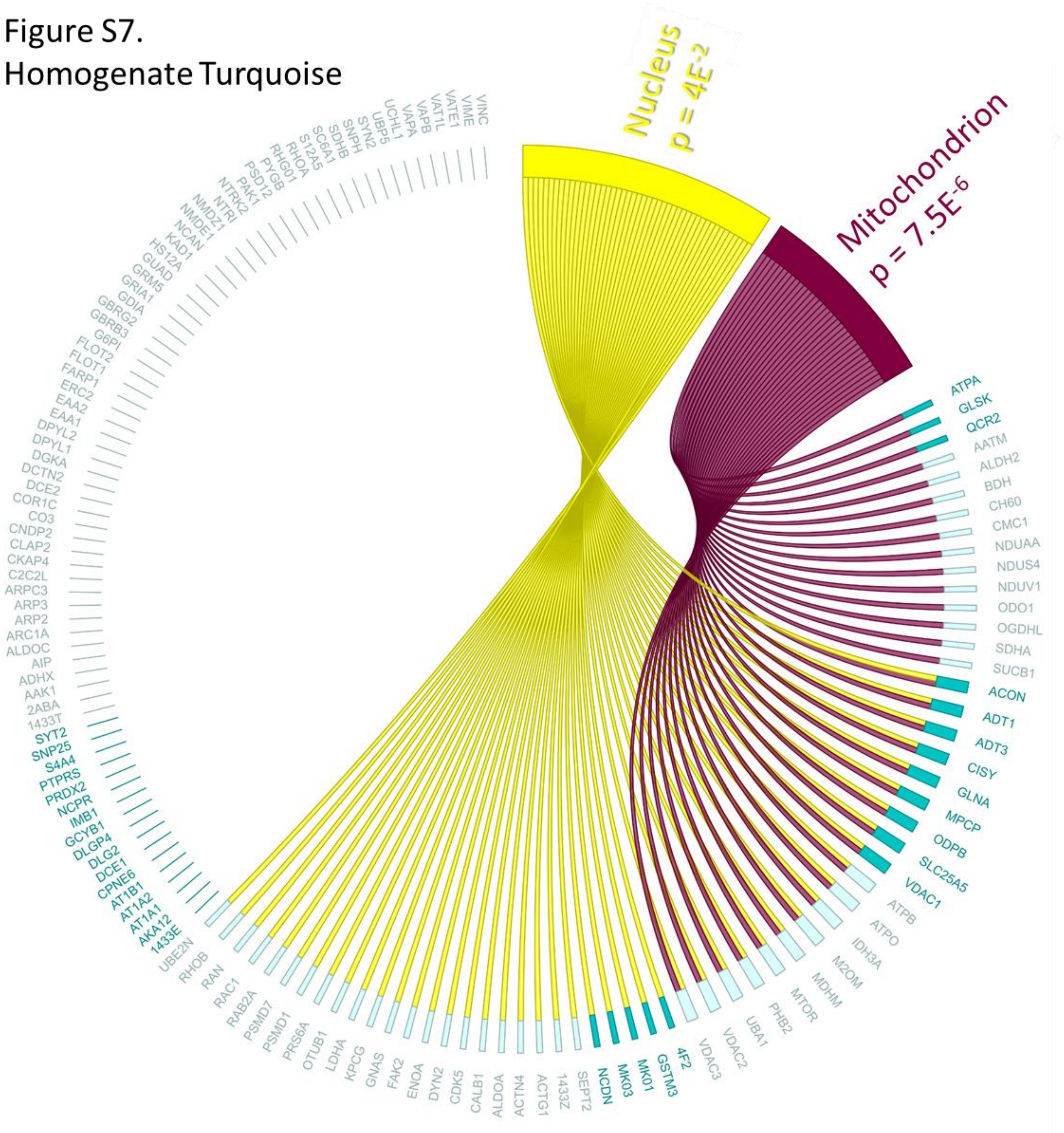

Figure S8.
Homogenate Orange

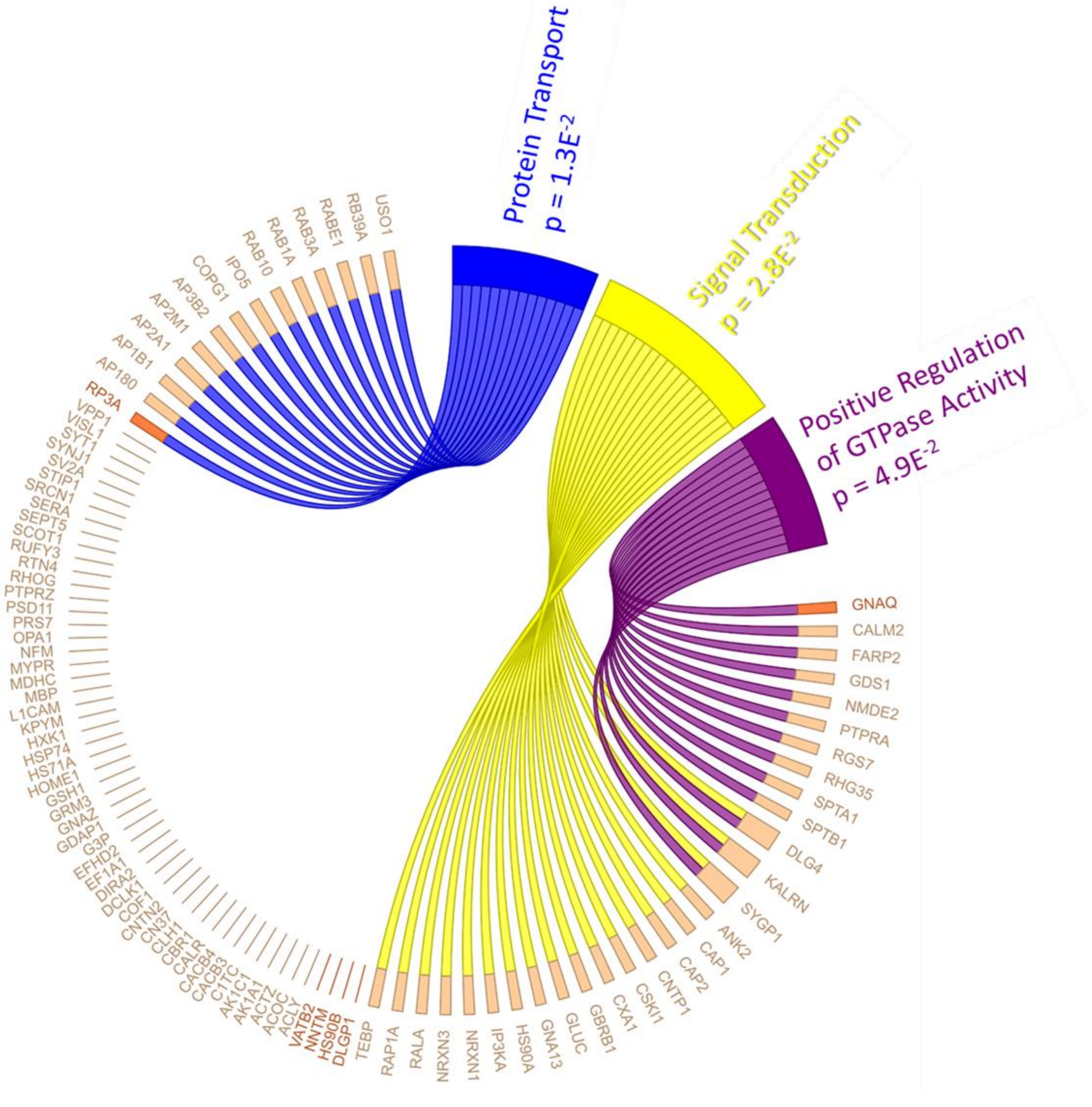

Figure S9.
Homogenate Green

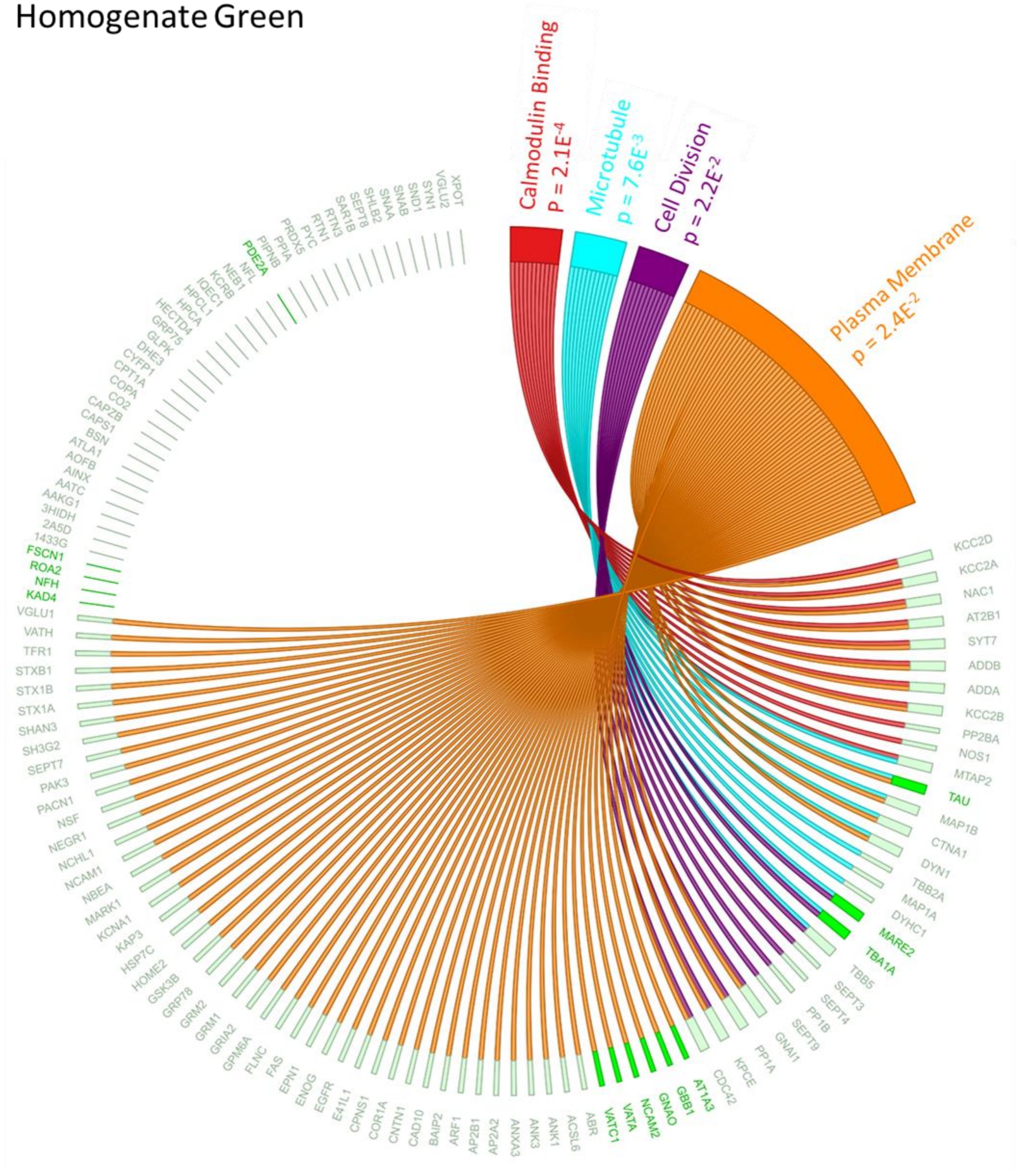

Figure S10.
Synaptosome Blue

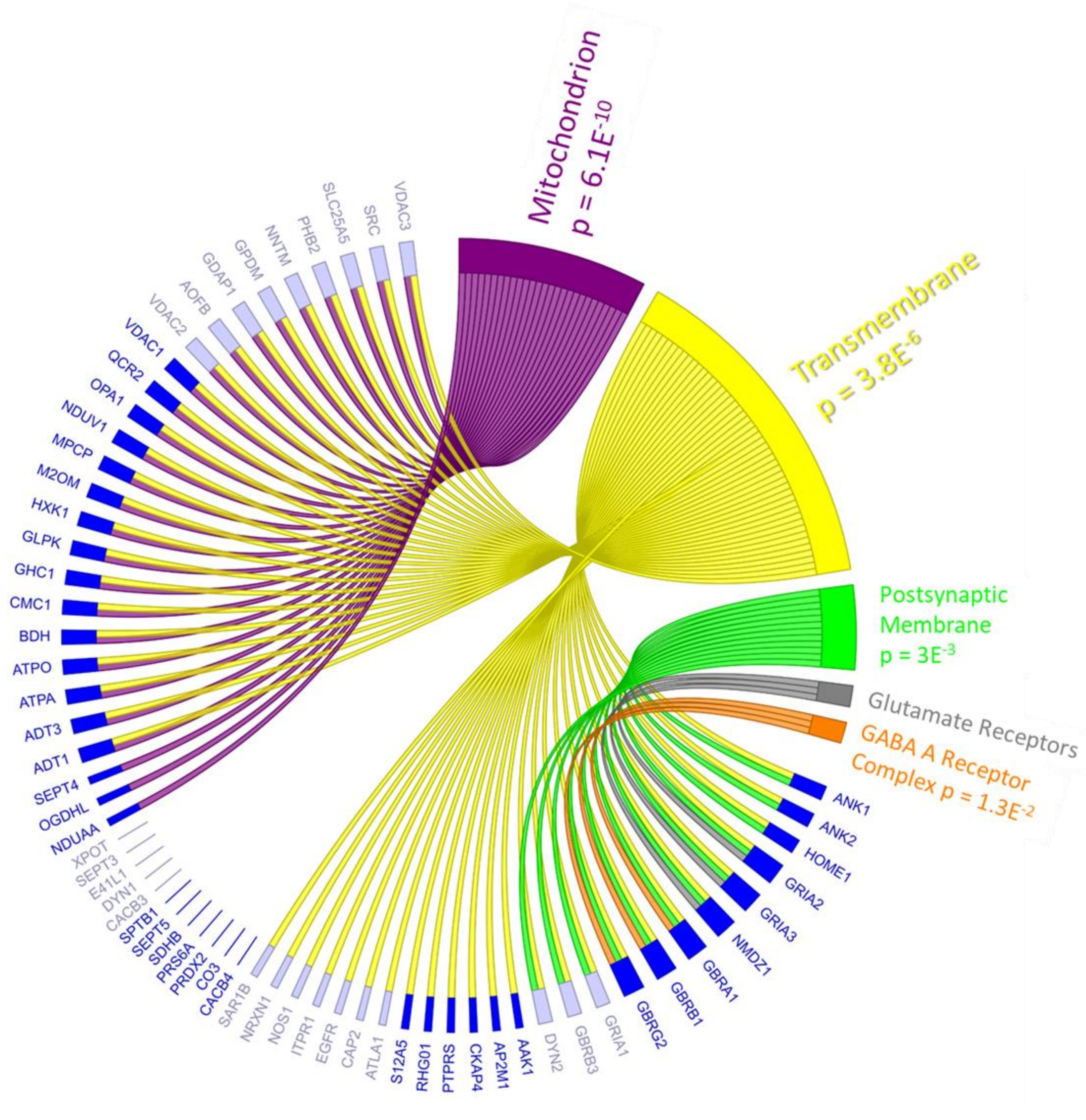

Figure S11.
Synaptosome Magenta

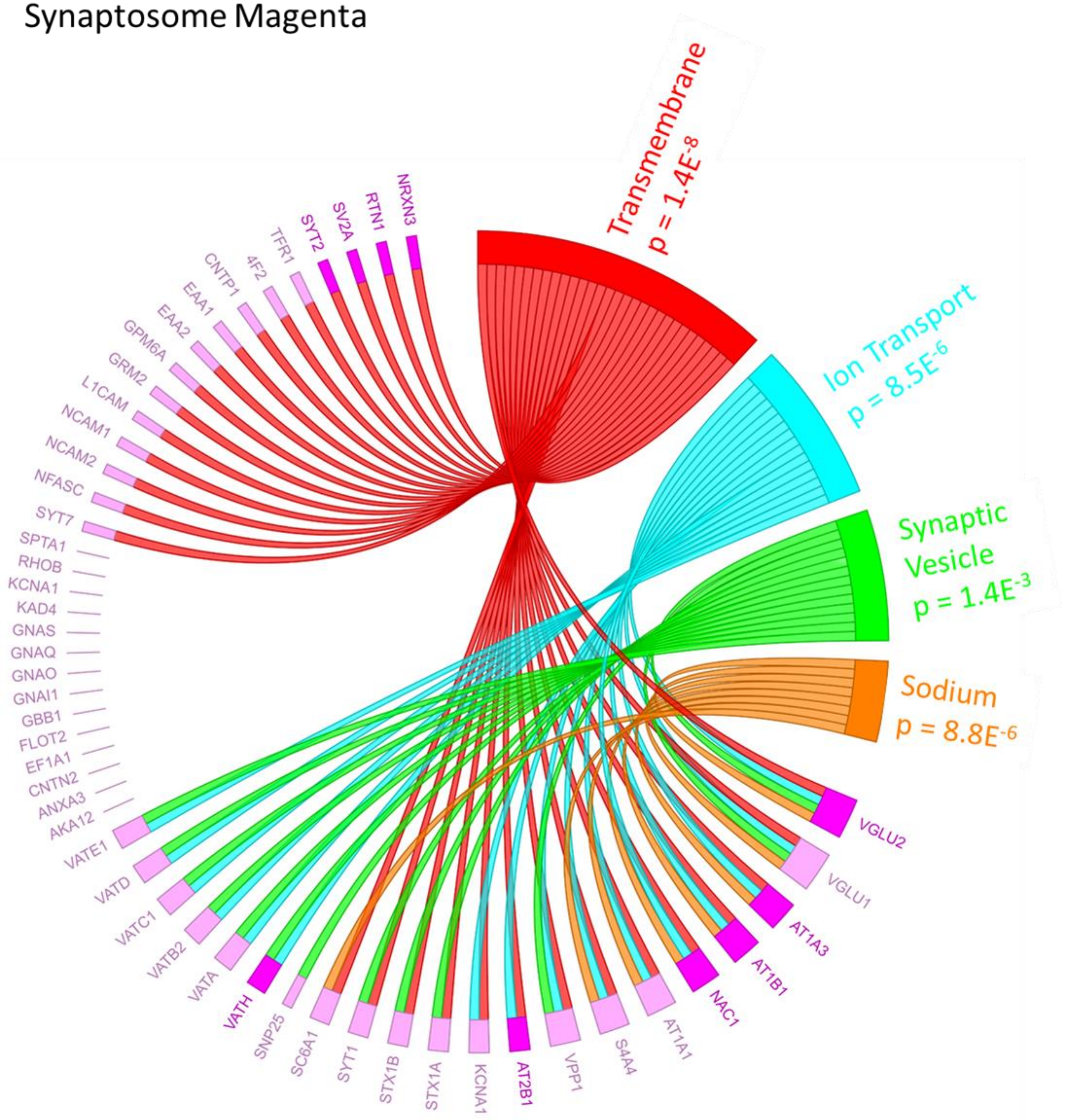

Figure S12.
Synaptosome Red

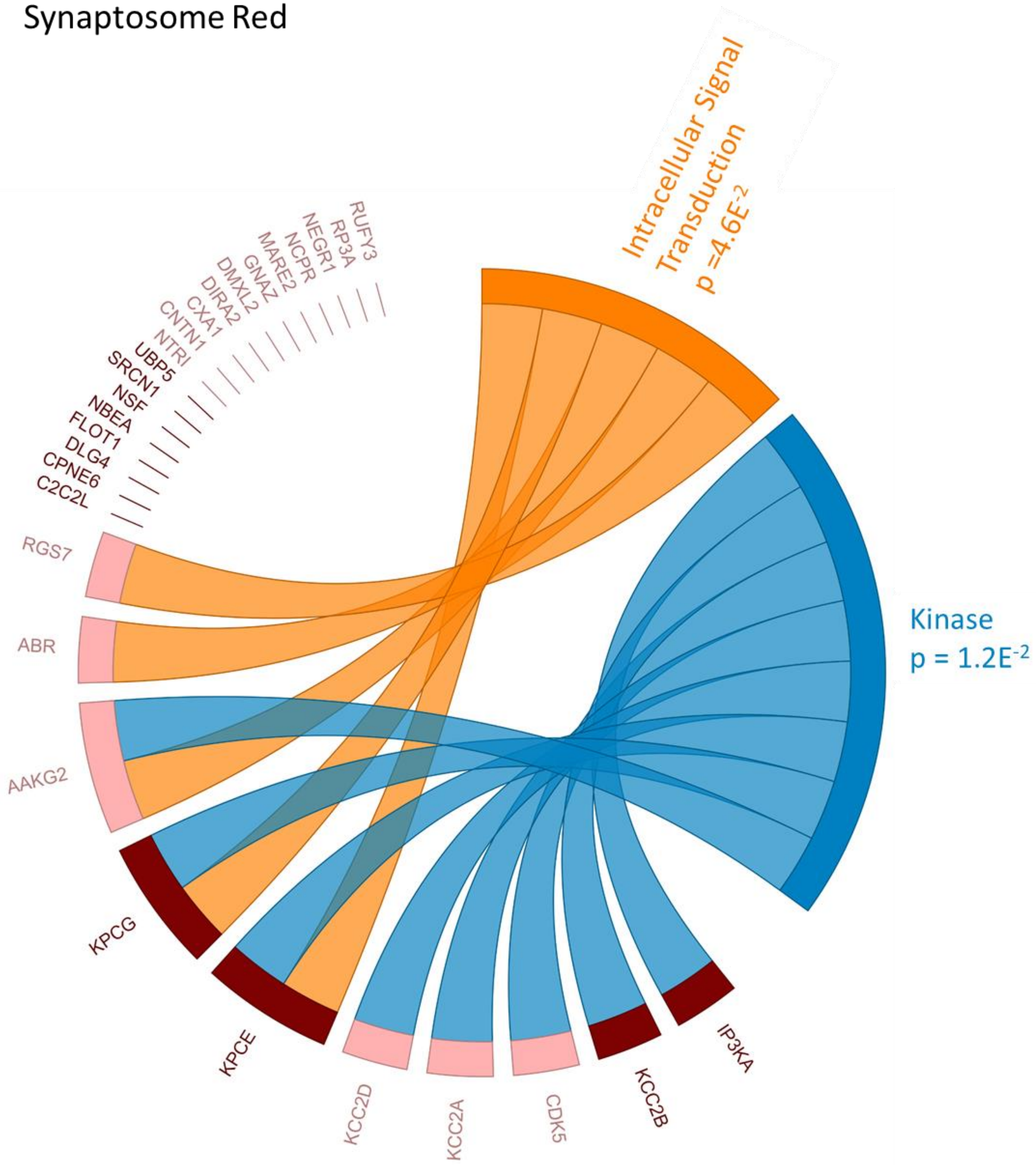

**Figure S13.** Associations between module Eigenprotein values. A. Reports the correlations between the module Eigenprotein values as a heat map. Thicker black border denotes a significant correlation ($p < 0.05$). B. Plots the Eigenprotein values of the homogenate Mitochondrion module (Turquoise) as well as the synaptosome Mitochondrion & Postsynaptic Membrane module (Blue), which are correlated in controls, but not Sz.

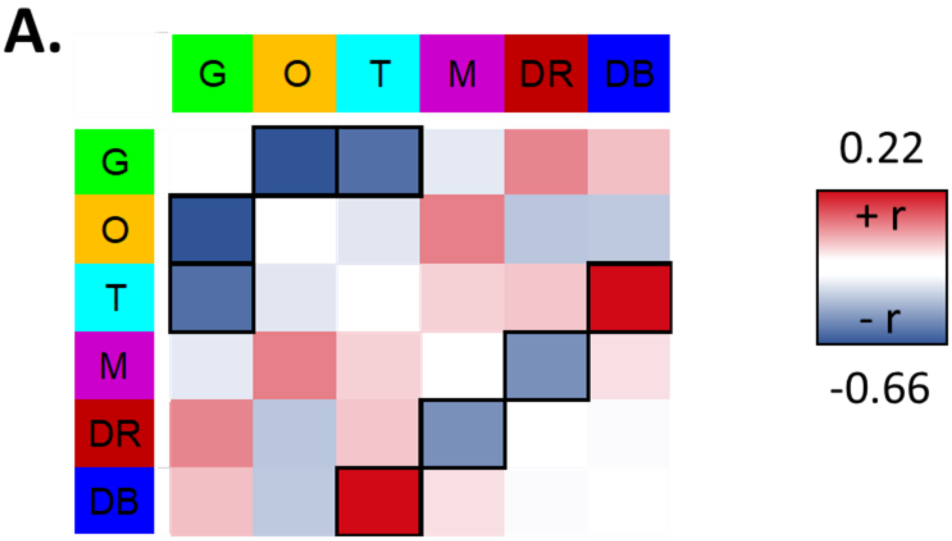


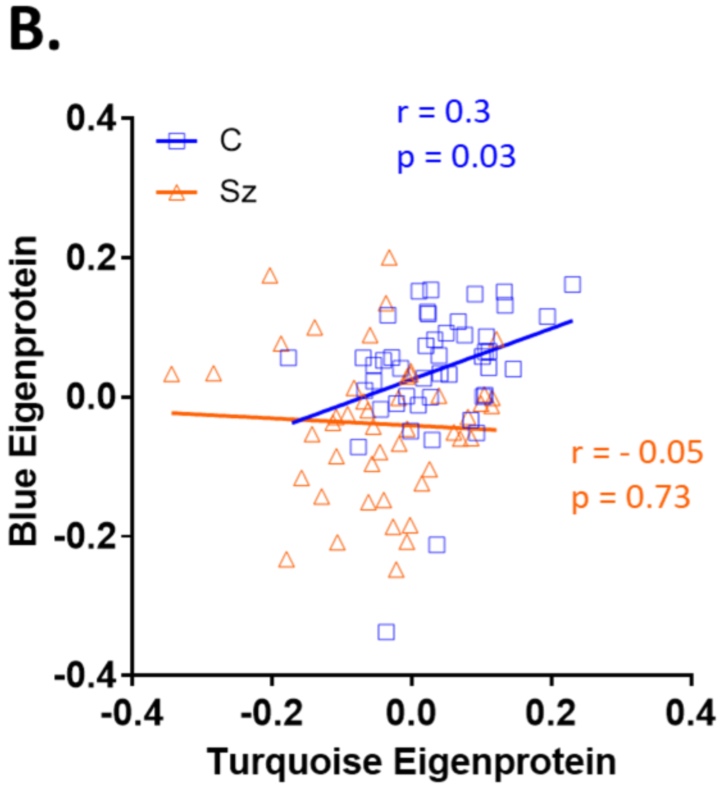

**Figure S13. Comparison of Proteomic and Transcriptomic Analysis of Sz tissue. A.** Chi-square analysis of bulk tissue RNAseq and homogenate protein level alterations. **B.** Chi-square analysis of bulk tissue RNAseq and synaptosome protein level alterations. **C.** Linear regression of bulk tissue RNAseq and homogenate protein level alterations. **D.** Linear regression of bulk tissue RNAseq and synaptosome protein level alterations.

**A.**

| Homogenate | Prot Up | Prot Down | Prot UnCh |
|---|---|---|---|
| mRNA Up | 7 | 6 | 74 |
| mRNA Down | 2 | 9 | 59 |
| mRNA UnCh | 11 | 20 | 212 |

chi-square p = 0.38

**B.**

| Synaptosome | Prot Up | Prot Down | Prot UnCh |
|---|---|---|---|
| mRNA Up | 1 | 7 | 23 |
| mRNA Down | 3 | 11 | 12 |
| mRNA UnCh | 5 | 37 | 55 |

chi-square p = 0.21

**C.**

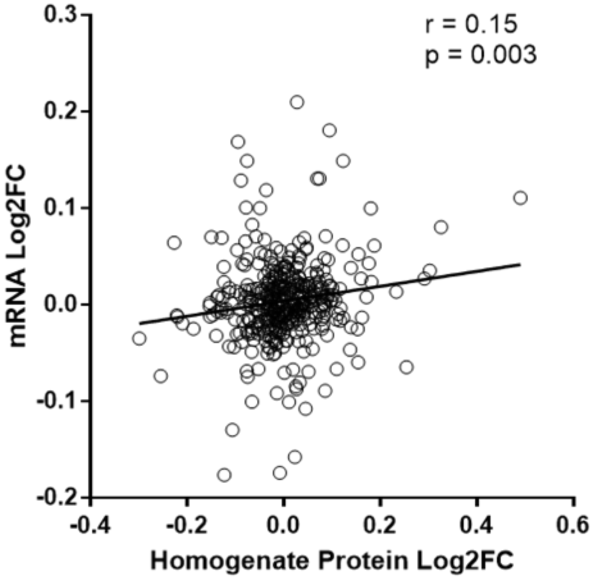


**D.**

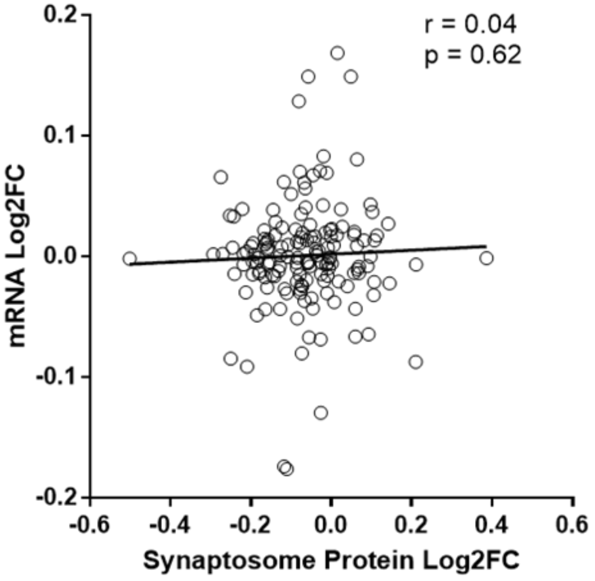

**Figure S14. Comparison of Limma-Voom p-values with and without pH as a covariate.** Unadjusted p-values for the Sz-control comparison, with and without pH as a covariate, are plotted for the homogenate (**A.**) and Synaptosome (**B.**) preparations. While there was a uniform shift in p-values with the inclusion of pH, the rank ordering of the effects were largely unaltered.

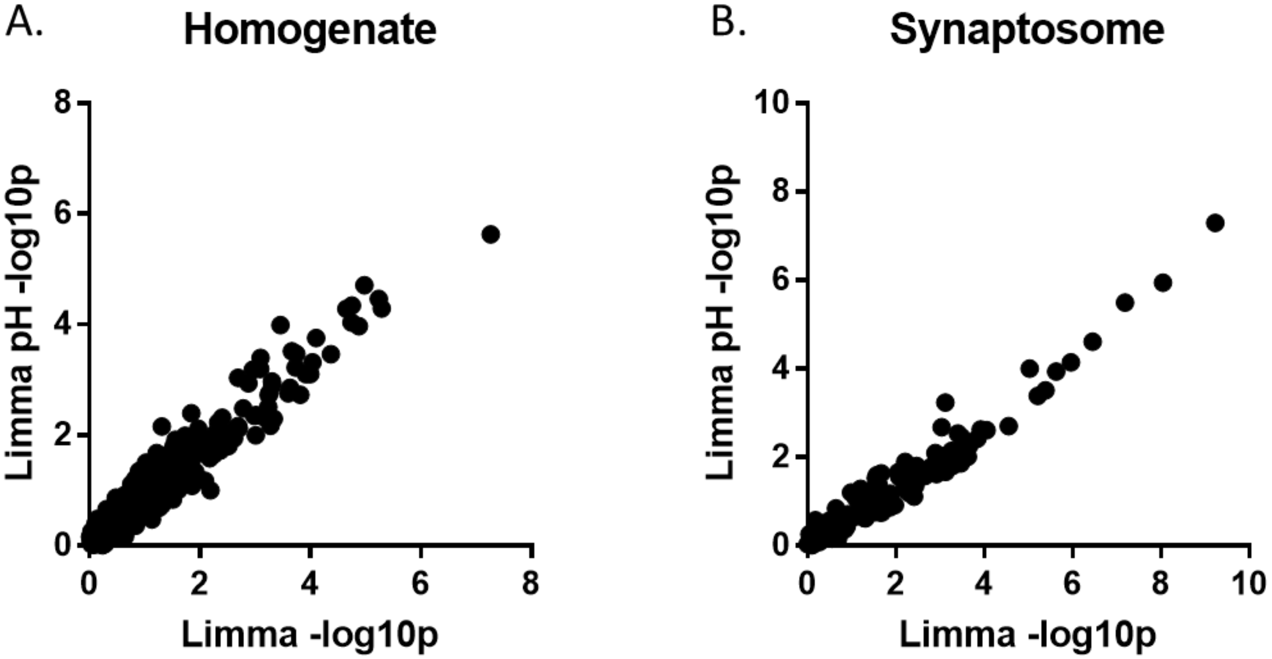
